# Bio-hybrid micro-swimmers propelled by flagella isolated from C. reinhardtii^†^

**DOI:** 10.1101/2021.05.19.444861

**Authors:** Raheel Ahmad, Albert J Bae, Yu-Jung Su, Samira Goli Pozveh, Eberhard Bodenschatz, Alain Pumir, Azam Gholami

**Affiliations:** Max Planck Institute for Dynamics and Self-Organization, Am Fassberg 17, D-37077 Göttingen, Germany; Lewis & Clark College, Portland, Oregon, USA; Institute for Dynamics of Complex Systems, University of Göttingen, Göttingen 37077, Germany; Laboratory of Atomic and Solid-State Physics and Sibley School of Mechanical and Aerospace Engineering, Cornell University, Ithaca, New York 14853, USA; Laboratoire de Physique, Ecole Normale Supérieure de Lyon, Université Lyon 1 and CNRS, F-69007 Lyon, France

## Abstract

Bio-hybrid micro-swimmers, composed of biological entities integrated with synthetic constructs, actively transport cargo by converting chemical energy into mechanical work in a fluid at low Reynolds number, where viscous drag dominates over inertia. Here, using isolated and demembranated flagella from green algae *Chlamydomonas reinhardtii* (*C. reinhardtii*), we build efficient axonemally-driven micro-swimmers that consume ATP to propel micron-sized beads. Depending on the calcium concentration, we observed two main classes of motion: Whereas beads move along curved trajectories at calcium concentrations below 0.03 mM, they are propelled along straight paths when the calcium concentration increases. In this regime, they reached velocities of approximately 20 *μ*m/sec, comparable to human sperm velocity *in vivo*. We relate this transition to the properties of beating axonemes, in particular the reduced static curvature with increasing calcium concentration. To quantify the motion, we used mode decomposition of the flagellar waveform, and we studied both analytically and numerically the propulsion of the bead as a function of the axonemal waveform and bead-axoneme attachment geometry. While our analysis semi-quantitatively describes the experimental results, it also reveals the existence of a counter-intuitive propulsion regime where the speed of the axonemally-driven bead increases with the size of the bead. Moreover, we demonstrated that asymmetric, sideways attachment of the axoneme to the bead can also contribute to the rotational velocity of the micro-swimmer. The uncovered mechanism has potential applications in the fabrication of synthetic micro-swimmers, and in particular, bio-actuated medical micro-robots for targeted drug delivery.

## 1 Introduction

Bio-actuated micro-swimmers, fabricated by integration of a live biological entity with a synthetic cargo, are the subject of growing interest, in particular for their potential applications in the field of targeted drug delivery or assisted fertilization *in vivo* ^1–3^. It is known that an essential aspect in designing bio-hybrid micro-swimmers is to take advantage of biological motors, which are highly efficient in converting chemical energy into mechanical work^4^. Therefore, the main goal in the synthesis of bio-hybrid micro-swimmers is to integrate these highly efficient energy conversion modules into bio-compatible nano- and micro-swimmers to realize various tasks. In the past decade, the sensory capabilities and the autonomous motility and power of various bacterial species, mainly *E. coli* have been extensively utilized as a biological micro-actuator in bio-hybrid micro-swimmers to provide an efficient cargo (e.g. liposomes or coated beads) transport^5–12^.

A wide variety of motility mechanisms exist in nature. This includes the polarized assembly and disassembly of biopolymers for the directional movement of amoeboid cells ^13,14^, or motility driven by eyelash-shaped cellular protrusions - cilia and flagella - that perform periodic whip-like motion to generate fluid flow and propulsion^15–17^. In many organisms, multiple motile cilia coordinate their beating activity to enhance the swimming efficiency or directional fluid transport. Examples are transport of cerebrospinal fluid, which contains physiologically important signaling molecules, through the ventricular cavities of the mammalian’s brain^18^, mucus clearance in mammalian airways^19^, or the bi-flagellated algae *Chlamydomonas reinhardtii* (*C. reinhardtii*) which swims in water with its two flagella using a characteristic breaststroke beat pattern ^20–24^. Cilia and flagella have a tube-shaped structure consisting of a periodic arrangement of nine microtubule doublets (MTDs) at the periphery and two axial singlet microtubules at the center. This 9+2 structure, known as axoneme, has the diameter of 200 nm and can be actuated via co-ordinated collective activity of dynein molecular motors that convert chemical energy released during ATP hydrolysis into mechanical work by sliding neighboring MTDs^25–33^. Flagella in different species show a variety of beat patterns. In contrast to spermatozoa (typical length 50 *μ*m), flagella isolated from green algae *C. reinhardtii* (typical length 10 *μ*m) beat by asymmetric bending deformations propagating at a fundamental beat frequency along the axonemal length in a base-to-tip direction ^34–38^. The observed asymmetric waveform is such that the flagellar shapes over one beat cycle average to a semi-circular arc with a mean curvature of about −0.2 *μ*m^−1^. This static component of the axonemal curvature leads to a curved swimming trajectory ^36,39–41^, whereas in the absence of this component, the flagella swims along a straight path^34,35^.

In the present work, we demonstrate the feasibility of using isolated and demembranated flagella from *C. reinhardtii* as a bioactuator to carry and maneuver micron-sized particles. To the best of our knowledge, there has been no report in the literature on isolated flagella-driven propulsion of cargo. Figure 1 demonstrates the existence of two different regimes of propulsion. At low calcium concentration, see Fig. 1A-B, the micro-swimmer is propelled on a highly curved trajectory. At higher calcium concentration, on the other hand, the bead is propelled along an almost straight path, at a velocity of ~ 20 *μ*m/s, comparable to the velocity of human sperm *in vivo*, see Fig. 1C-D. The possibility to externally modify flagellar wave components and beating frequency with light or chemical stimuli (such as calcium ions, as in Fig. 1B,D) ^35,42–44^ opens interesting perspective to control such a bio-hybrid micro-swimmer. The micro-swimmers, propelled by isolated and reactivated flagella, are potentially attractive to develop minimally invasive devices for medical applications, such as *in vivo* active cargo transport^7,8^. In this context, understanding the propulsion mechanism as a function of the flagellar beat pattern and flagella-bead attachment geometry appears as an important pre-requisite.

**Fig. 1.**
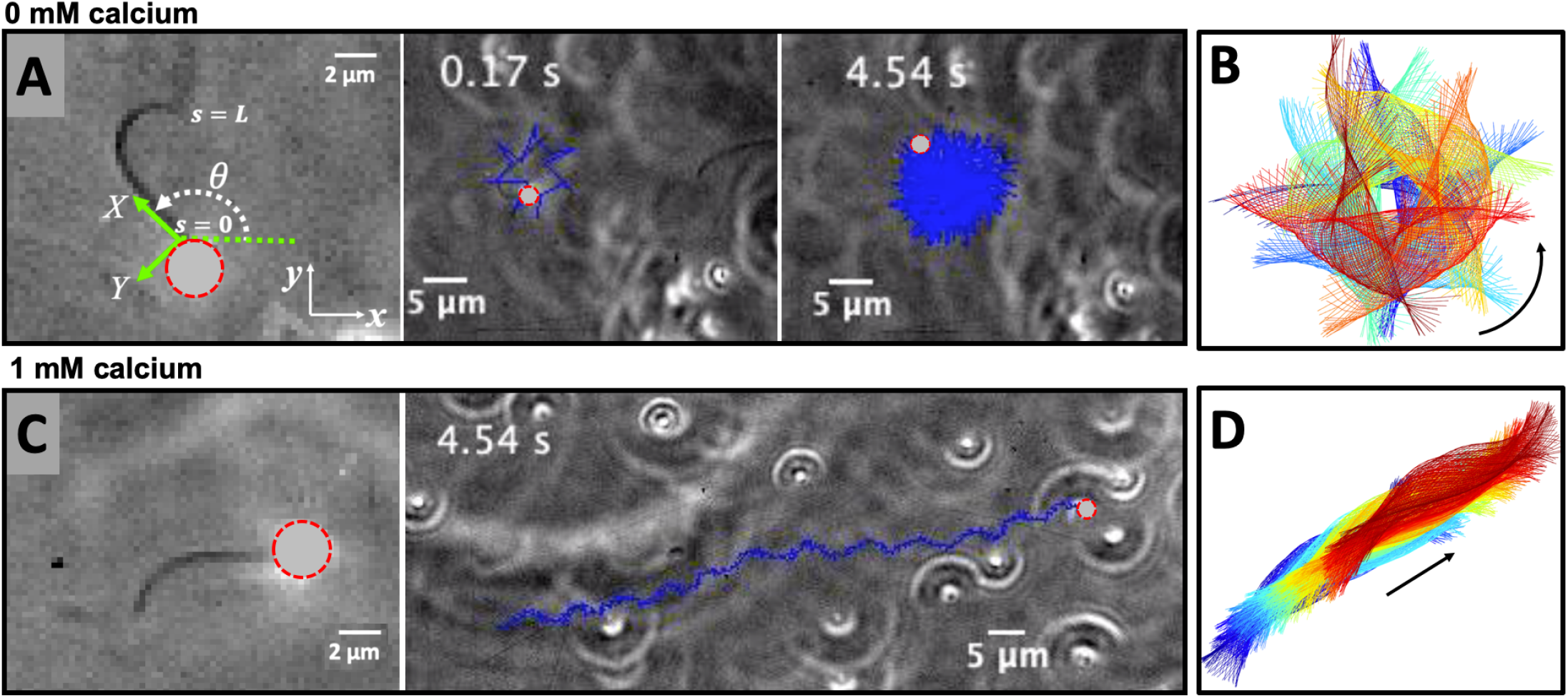
Axonemally-driven beads and the effect of calcium ions. A) An exemplary axoneme at [Ca^2+^]=0 mM is attached at the basal end (*s* = 0) to a bead of diameter 1 *μ*m. The blue trajectory of the bead center shows a curved swimming path at two different time points. B) The time-projection of the axonemal shapes highlights the counter-clockwise rotation of the axoneme as bending deformations propagate along the contour length. C) In the presence of 1 mM calcium, the bead is propelled on average along a straight path at the propulsion speed of around 20 *μ*m/sec, which is comparable to the human sperm propulsion speed in mucus. D) Calcium ions trigger a transition from an ‘asymmetric’ flagellar waveform that rotates counter-clockwise to a ‘symmetric’ waveform that swims along a straight path. In both experiments, the ATP concentration is 200 *μ*M (see Videos 1,2).

Here, we combined experiments, numerical simulations and analytical approximations to investigate the effect of (i) external calcium ions, (ii) various wave components, (iii) the cargo size, and (iv) symmetric versus asymmetric bead-axoneme attachment on the swimming dynamics of a bead that is propelled by an ATP-reactivated axoneme. We used calcium ions to reduce the static curvature of axonemes by one order of magnitude, thereby^34,35^ inducing a transition from circular to straight swimming trajectories of axonemal-propelled beads (see Fig. 1). Furthermore, we described the axonemal waveform as a combination of a static curvature and a traveling wave component and used resistive-force theory ^45,46^ to obtain analytical approximations for the translational and rotational velocities of an axonemal-propelled bead in the limit of small amplitude of curvature waves, and simulated the swimming trajectories. Our analysis revealed a surprising non-monotonic behavior of the mean translational and rotational velocities of the axonemal-driven bead as a function of the bead radius. Finally, we showed that for a freely swimming axoneme, which rotates predominantly with its static component of the axonemal waveform, sideways bead attachment is sufficient to generate mean rotational velocities comparable to the rotation rates induced by the static curvature.

## 2 Materials and Methods

### 2.1 Isolation of axonemes from *C. reinhardtii* and attachment to the beads

Isolation and reactivation of axonemes are done based on a standard protocol described in Refs. 47,48. Briefly, wild-type *C. reinhardtii* cells, strain SAG 11-32b, are grown axenically in TAP (trisacetate-phosphate) medium on a 12 h/12 h day-night cycle. Upon adding Dibucaine, cells release their flagella which we purify on a 25% sucrose cushion, and demembranate them using detergent NP-40 in HMDEK solution (30 mM HEPES-KOH, 5 mM MgSO_4_, 1 mM DTT, 1 mM EGTA, 50 mM potassium acetate, pH 7.4) supplemented with 0.2 mM Pefabloc. The membrane-free axonemes are resuspended in HMDEK buffer plus 1% (w/v) polyethylene glycol (M_*w*_ = 20 kgmol^−1^), 0.2 mM Pefabloc and used freshly after isolation. To perform reactivation experiments, we diluted the axonemes in HMDEKP reactivation buffer (HMDEK plus 1 % PEG) supplemented with ATP at different concentrations. Beads (Polystyrene microspehere) were washed twice with HMDEKP buffer supplemented with 0.01% SDS and resuspended in the same solution, and sonicated for 5 min before use. For bead attachment, beads are incubated for 20 min with axonemes before being mixed with reactivation buffer and infused into 100 *μ*m deep flow chambers, built from cleaned glass and double-sided tape. The glass slides are treated with casein (2 mg/mL in HMDEKP buffer) for 5 min before use to avoid axoneme-substrate attachment^38^. For calcium experiments, we added CaCl_2_ at different concentrations to the HMDEKP reactivation buffer.

### 2.2 Axoneme Tracking

We used high-speed phase-contrast microscopy (100X objective, 1000 fps) to capture high-resolution images of fast beating axonemes that propel micron-sized beads and swim parallel to the glass substrate with an approximately planar axonemal beat. This effective confinement to 2D significantly facilitates tracking of flagellar shapes and their analysis. In these images, the bead appears as a bright spot, which interferes with the tracking algorithm of the axoneme. Therefore, the first step is to filter out the bead by applying a threshold to the images (Fig. S1). Next, to increase the signal to noise ratio, we invert the phase-contrast images and subtract the mean-intensity of the time series ^38^. To smooth the images, we apply a Gaussian filter. Tracking of axonemes is done using a customized MATLAB code (available in the supplemental information) which is based on the gradient vector flow (GVF) technique^49–51^. In this algorithm, for the first frame, we select a region of interest that includes only one actively beating axoneme. Then, to initialize the snake, we draw a line polygon along the contour of the axoneme in the first frame (see Fig. S1). This polygon is interpolated at *N* equally spaced points and used as a starting parameter for the snake. The GVF is calculated using the GVF regularization coefficient *μ* = 0.1 with 20 iterations. The snake is then deformed according to the GVF where we have adapted the original algorithm by Xu and Prince for open boundary conditions^49,50^. The axoneme is characterized by *N* points along the axonemal length *s* so that *s* = 0 is the basal region and *s* = *L* is the distal tip. Here *L* is the total contour length of the filament. The position of axoneme at contour length *s*_*i*_ and time point *t* is given by **r**(*s*_*i*_, *t*) = (*x*(*s*_*i*_, *t*), *y*(*s*_*i*_, *t*)). Furthermore, to estimate the systematic error of our tracking algorithm, we generated artificial filaments with known values of mean curvature and used the GVF algorithm to determine the curvature from the tracking algorithm. As shown in Fig. S2, the measured values systematically deviate from the real values at small mean curvatures, but provide a much better estimate when the mean curvature is larger. In other words, our algorithm has a smaller systematic error (less than 4%) for curved filaments with dimensionless mean curvature *C*_0_ larger than 0.3.

### 2.3 Resistive-force theory and calculations of mean translational and rotational velocities

The fluid flow generated by the swimming of small objects is characterized by very small Reynolds numbers. In this regime, viscous forces dominate over inertia and non-reciprocal motion is necessary to break the time-symmetry and generate propulsion (scallop theorem) ^52,53^. The micro-swimmer in our system is formed of an axoneme (a filament of characteristic length *L* ~ 10 *μ*m and radius 0.1 *μ*m), which is attached from one end to a micron-sized bead and swims in an aqueous solution of viscosity *μ* = 10^−3^ Pa s and density *ρ* = 10^3^ kg m^−3^. Given the characteristic axonemal wave velocity *V* = *λ/T* ≈ 0.5 mm s^−1^ (calculated for a typical axonemal beat frequency of 50 Hz and a wavelength which is comparable to the axonemal contour length *L*), the Reynolds number *Re* = *ρLV/μ* will be a small number around 0.005. An important feature of swimming at low Reynolds numbers is its reversibility, which is a consequence of the linearity of the Stokes equation ^53^. In this physical regime, Newton’s laws then consist of an instantaneous balance between external and fluid forces and torques exerted on the swimmer, i.e. **F**_ext_ + **F**_fluid_ = 0 and ***τ***_ext_ + ***τ***_fluid_ = 0. The force **F**_fluid_ and torque ***τ***_fluid_ exerted by the fluid on the axoneme-bead swimmer can be written as

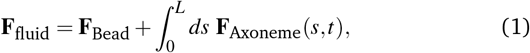

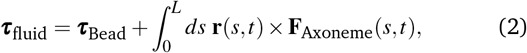

where **F**_bead_ and ***τ***_Bead_ are the hydrodynamic drag force and torque acting on the bead, and the integrals over the contour length *L* of the axoneme calculate the total hydrodynamic force and torque exerted by the fluid on the axoneme. The bead is propelled by the oscillatory shape deformations of an ATP-reactivated axoneme. At any given time, we consider axoneme-bead swimmer as a solid body with unknown translational and rotational velocities **U**(*t*) and **Ω**(*t*) yet to be determined. **F**_fluid_ and ***τ***_fluid_ can be separated into propulsive part due to the relative shape deformations of the axoneme in the body-fixed frame and the drag part ^54^

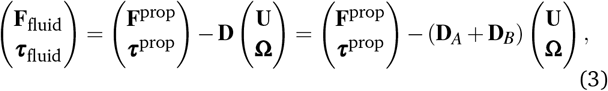

where the 6*×*6 geometry-dependent drag matrix **D** is symmetric and non-singular (invertible) and is composed of drag matrix of the axoneme **D**_*A*_ and drag matrix of the bead **D**_*B*_. We also note that a freely swimming axoneme-bead experiences no external forces and torques, thus **F**_fluid_ and ***τ***_fluid_ must vanish. Further, since swimming effectively occurs in 2D, **D** is reduced to a 3*×*3 matrix and Eq. 3 can be reformulated as

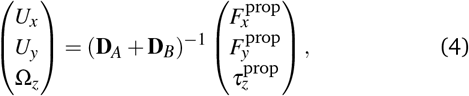

which we use to calculate translational and rotational velocities of the swimmer after determining the drag matrices **D**_*A*_ and **D**_*B*_, and the propulsive forces and torque 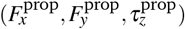.

We calculate 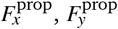 and 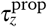 in the body-fixed frame by selecting the basal end of the axoneme (bead-axoneme contact point) as the origin of the swimmer-fixed frame. As shown in Fig. 1A and Fig. 2A, we define the local tangent vector at contour length *s* = 0 as **X**-direction, the local normal vector ***n*** as the **Y**-direction, and assume that ***z*** and **Z** are parallel. Here (***x***,***y***,***z***) denote an orthogonal lab-frame basis. We define *θ*_0_(*t*) = *θ* (*s* = 0, *t*) as the angle between ***x*** and **X** which gives the velocity of the bead in the laboratory frame as 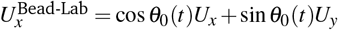 and 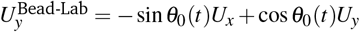. Furthermore, note that the instantaneous velocity of the axoneme in the lab frame is given by **u** = **U** + **Ω** *×* **r**(*s, t*) + **u**′, where **u**′ is the deformation velocity of the flagellum in the body-fixed frame, **U** = (*U*_*x*_,*U*_*y*_, 0) and **Ω** = (0, 0, Ω_*z*_) with Ω_*z*_ = *dθ*_0_(*t*)*/dt*.

**Fig. 2.**
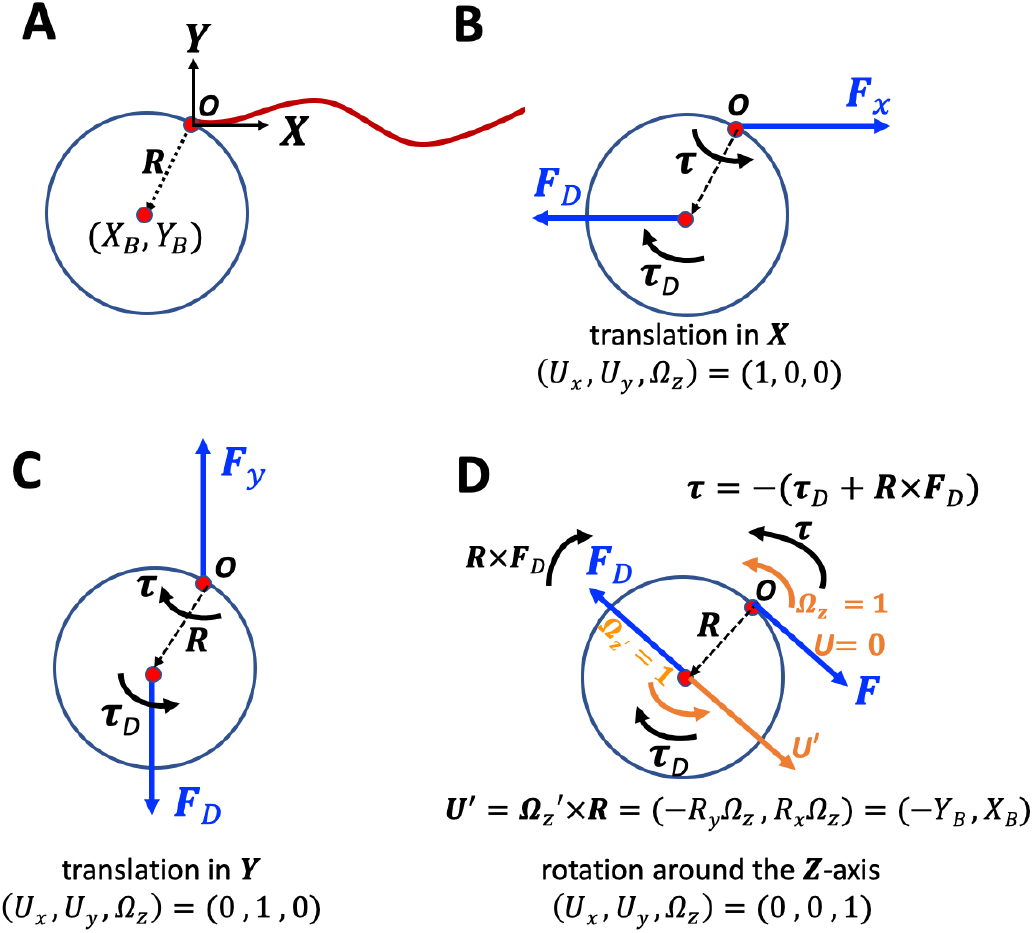
A) Illustration of the bead orientation with respect to the axoneme and definition of the swimmer-fixed frame. Tangent vector at *s* = 0 (basal end) defines the **X**-direction. B-D) Schematic drawing of the forces and torques that counteract the hydrodynamic drag force and torque.

To calculate 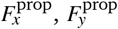 and 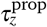 for a given beating pattern of axoneme in the body-fixed frame, we used the classical framework of resistive-force theory (RFT), which neglects long-range hydrodynamic interactions between different parts of the flagellum as well as the inter-flagella interactions ^45,46^. In this theory, the flagellum is discretized as a set of small rode-like segments moving with velocity **u**′(*s, t*) in the body-frame and the propulsive force **F**^prop^ is proportional to the local center-line velocity components of each segment in parallel and perpendicular directions

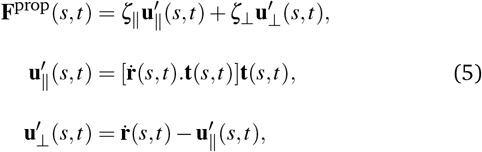

where 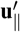 and 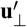 are the projections of the local velocity on the directions parallel and perpendicular to the axoneme. The friction coefficients in parallel and perpendicular directions are defined as *ζ*_‖_ = 4*πμ/*(ln(2*L/a*) + 0.5) and *ζ*_⊥_ = 2*ζ*_‖_, respectively. This anisotropic hydrodynamic friction experienced by a cylindrical segment is the basis of propulsion by a flagellum. For a water-like environment with dynamic viscosity *μ* = 10^−3^ Pa s, we obtain *ζ*_‖_ ~ 2.1 pN msec*/μ*m^2^. Notably, the resulting velocity is not parallel to the propulsive force **F**^prop^.

Here is a brief summary of the steps in the RFT analysis: First, we translate and rotate the axoneme such that the basal end is at position (0, 0) and the local tangent vector at *s* = 0 at any time *t* is along the ***x***-axis (see Fig. 1A). In this way, we lose the orientation information of the axoneme at all the time points except for the initial configuration at time *t* = 0. Second, we calculate propulsive forces and torque in the body-frame using RFT (Eq. 5), and then use Eq. 4 to obtain translational velocities *U*_*x*_, *U*_*y*_ as well as rotational velocity Ω_*z*_ of the axoneme. Now the infinitesimal rotational matrix can be expressed as

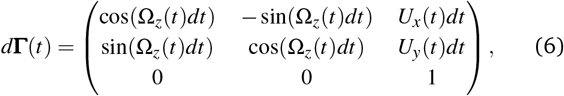

which we use to update rotation matrix as **Γ**(*t* + *dt*) = **Γ**(*t*)*d* **Γ**(*t*), considering **Γ**(*t* = 0) to be the unity matrix. Having the rotation matrix at time *t*, we obtain the configuration of the axoneme at time *t* from its shape at the body-fixed frame by multiplying the rotation matrix as **r**_lab-frame_(*s, t*) = **Γ**(*t*)**r**_body-frame_(*s, t*), which can then be compared with experimental data. Please note that **r**_lab-frame_(*s, t*) is an input from experimental data presenting the beating patterns in the lab frame.

#### 2.3.1 Drag matrix of a bead in 2D

Let us consider the two-dimensional geometry defined in Fig. 2A. Note that the origin of the swimmer-fixed frame is not at the bead center; rather it is selected to be at the bead-axoneme contact point, as shown in Fig. 2A. In general, the tangent vector at position *s* = 0 of the axoneme, which defines the **X**-axis, does not pass through the bead center located at (*X*_*B*_,*Y*_*B*_). This asymmetric bead-axoneme attachment is also observed in our experiments, as shown in Fig. 1A. We emphasize that the drag force is actually a distributed force, given by *d***f** = *σ*.*d***A**, applied at the surface of the sphere, but symmetry implies that drag force effectively acts on the bead center. We define the translational and rotational friction coefficients of the bead as *ν*_*T*_ = 6*πα*_*t*_ *μR* and *ν*_*R*_ = 8*πα*_*r*_*μR*^3^, where *μ* = 10^−3^ Pa s is the dynamic viscosity of water and factors *α*_*t*_ = 1*/* 1 − 9(*R/h*)*/*16 + (*R/h*)^3^*/*8 and *α*_*r*_ = 1*/* 1 − (*R/h*)^3^*/*8 are corrections due to the fact that axonemal-based bead propulsion occurs in the vicinity of a substrate ^55^. Here *R* is the bead radius (~ 0.5 *μ*m), *h* is the distance between the center of the bead and the substrate. Assuming *R/h* ~ 1, we obtain *α*_*t*_ = 16*/*9 and *α*_*r*_ = 8*/*7. We now look at each component of velocity and ask what force do we need to apply to counteract the viscous force and torque?

i. Translation in **X**-direction. In this case, we have (*U*_*x*_,*U*_*y*_, Ω_*z*_) = (1, 0, 0) as shown in Fig. 2B. We need to apply a force in +**X**-direction to counteract the drag force as

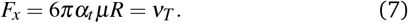

But we must also apply a torque in +**Z**-direction for the case illustrated in Fig. 2B (where *Y*_*B*_ *<* 0) to prevent rotation from occurring

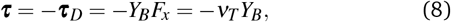

so we get

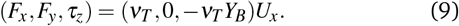
ii. Translation in **Y**-direction. This case corresponds to (*U*_*x*_,*U*_*y*_, Ω_*z*_) = (0, 1, 0) as shown in Fig. 2C. We have *F*_*x*_ = 0 and *F*_*y*_ = +6*πα*_*t*_ *μR* = *ν*_*T*_. Note that we need to apply a negative torque, and since *X*_*B*_ *<* 0, we have *τ*_*z*_ = +*X*_*B*_*ν*_*T*_ which gives

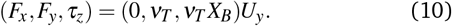
iii. Rotation around **Z**-direction. For this case, we have (*U*_*x*_,*U*_*y*_, Ω_*z*_) = (0, 0, 1) as shown in Fig. 2D. Before looking at the forces, let us examine the motion. The rotation 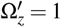 of the bead center around the origin *O* also generates translational velocity **U**′ = (−*R*_*y*_, *R*_*x*_)Ω_*z*_ = (−*Y*_*B*_, *X*_*B*_). Note that for *Y*_*B*_ *<* 0 and *X*_*B*_ *<* 0, we get 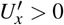 and 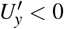 which is consistent. Around the center of the bead, drag exerts force and torque **F**_*D*_ and ***τ***_*D*_, as depicted in Fig. 2D

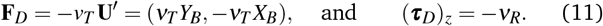

To counteract the drag force, we must apply

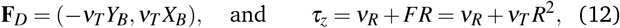

so we obtain (*F*_*x*_, *F*_*y*_, *τ*_*z*_) = (−*ν*_*T*_*Y*_*B*_, *ν*_*T*_ *X*_*B*_, *ν*_*R*_ + *ν*_*T*_ *R*^2^)Ω_*z*_.

Now we combine parts (i), (ii) and (iii) to obtain

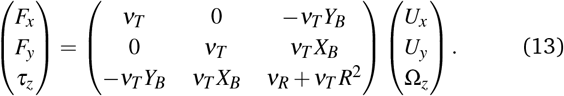

For the special case that center of the bead is at (*X*_*B*_,*Y*_*B*_) = (−*R*, 0) which corresponds to the situation that tangent vector of the flagella at *s* = 0 goes through the bead center, Eq. 13 simplifies to

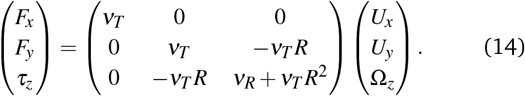

Note that Eqs. 13-14 present the forces and torque exerted by the bead on the fluid which has opposite sign of the forces generated by the fluid on the bead, so the drag matrix of the bead **D**_*B*_ is given by

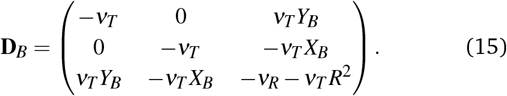

The general form of the drag matrix in 3D is derived in the next section.

#### 2.3.2 Drag matrix of a bead in 3D

Let us fix a couple of points *P* and *P*′ on or in a rigid body (see Fig. 3A). The distance between these points remains constant. Furthermore, if we attach two parallel vectors at *P* and *P*′, they would remain parallel under movement. Since this is the case, it follows that **Ω**_*P*_ = **Ω**_*P*′_, so we will drop the subscript and call it **Ω**.

**Fig. 3.**
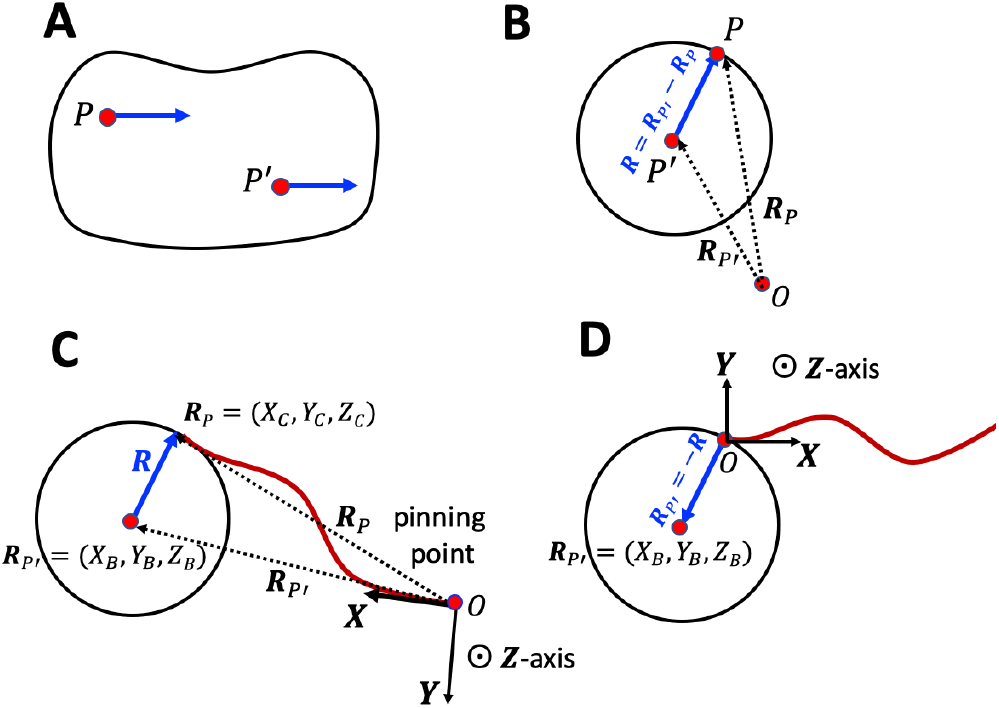
Schematic presentation of the set up and the coordinate system.

Let us denote the positions of *P* and *P*′ by **R**_*P*_ and **R**_*P*′_ (Fig. 3B). The distance |**R**_*P*′_ − **R**_*P*_| is constant, but the orientation changes

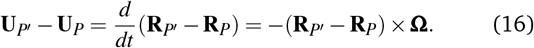

Next, suppose we apply a distribution of forces **f**_*i*_ at **R**_*i*_ as **F** = Σ_*i*_**f**_*i*_ which does not depend on *P* and *P*′. The torques depend on *P* and *P*′

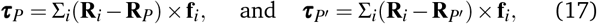

and the difference gives

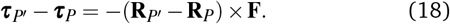

Now let us define the problem we wish to solve: What is the force **F** = (*F*_*x*_, *F*_*y*_, *F*_*z*_)^*T*^ and torque ***τ*** = (*τ*_*x*_, *τ*_*y*_, *τ*_*z*_)^*T*^ we need to apply at point *P* on the bead’s surface to make the bead move with translational velocity **U** = **U**_*P*_ and angular velocity **Ω**? This amounts to asking what is the drag force and drag torque one needs to overcome. Let us denote the center of the bead as *P*′ and define **R** = **R**_*P*_ − **R**_*P*′_ (Fig. 3B). This problem would have been much simpler if we were asked what force **F**′ and torque ***τ***′ needs to be applied at the bead’s center to counteract drag

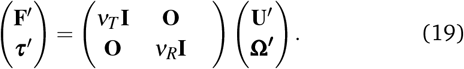

Here *ν*_*T*_ = 6*πRμ, ν*_*R*_ = 8*πR*^3^ *μ/*3 and

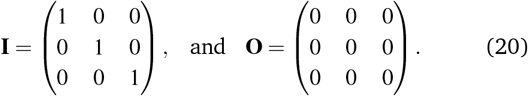

Now we use what we learned about rigid body motion in Eq. 16

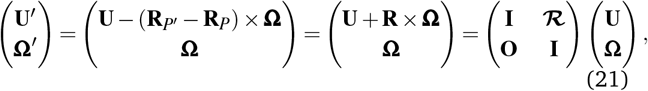

where we have defined

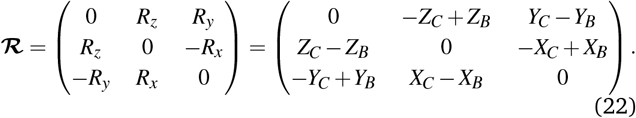

Recall that **R**_*P*_ = (*X*_*C*_,*Y*_*C*_, *Z*_*C*_) are the coordinates of the flagella-bead contact point and **R**_*P*′_ = (*X*_*B*_,*Y*_*B*_, *Z*_*B*_) are the coordinates of the bead center (Fig. 3C). Similarly, we have that

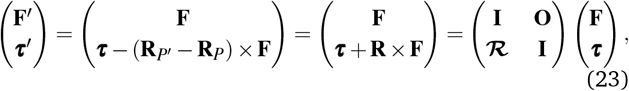

so the Eq. 19 becomes

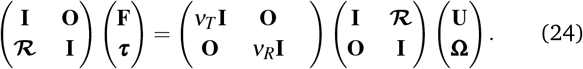

Multiplying both sides by 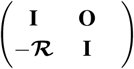 and calculating the products of matrices yields

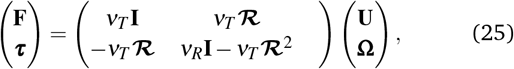

where

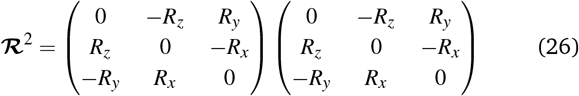

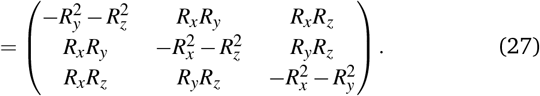

Possibly noteworthy is that

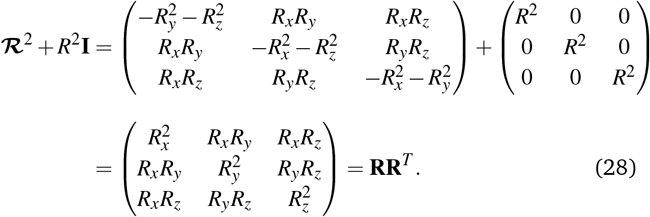

For the motion in *x* − *y* plane (2D), we are only interested in *F*_*x*_, *F*_*y*_ and *τ*_*z*_ as a function of *U*_*x*_, *U*_*y*_ and Ω_*z*_. Here we also assume *R*_*z*_ = 0, so we are only interested in components 1, 2, and 6

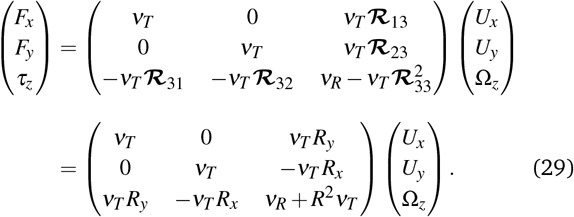

Up until now, we haven’t actually specified where our origin is. Let us set the origin to be at point *P* on the surface of the bead (**R**_*P*_ = 0). We will choose **X** to point tangent to the flagella at the point of attachment, and let the coordinates of the center of the bead be located at **R**_*P*′_ = (*X*_*B*_,*Y*_*B*_, 0). Note that *X*_*B*_ *≤* 0 and −*R ≤ Y*_*B*_ *≤ R* (Fig. 3D). Also note that **R** = −**R**_*P*′_, so the force equation reads

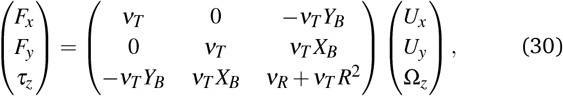

which was previously extracted in Eq. 13. Note that setting *Y*_*B*_ ≠ 0 allows us to handle the case where the flagella is not attached normal to the bead.

### 2.4 Principal component analysis

This analysis is based on the method introduced by Stephens et al. ^56^, to characterize the waveforms in *C. elegans*. We describe the shape of the flagellum by its unit tangent vector **t**(*s*) and the unit normal vector **n**(*s*) at distance *s* along the contour^57,58^. Instantaneous deformation of flagellum is described by curvature *κ*(*s, t*), using the Frenet-Serret formulas^56^

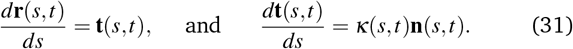

Let us define *θ* (*s, t*) to be the angle between the tangent vector at distance *s* and the ***x***-axis, then *κ*(*s, t*) = *dθ* (*s, t*)*/ds*. For shape analysis, we translate and rotate each flagellum such that basal end is at (0, 0) and the orientation of the tangent vector at *s* = 0 is along the ***x***-axis i.e. *θ* (*s* = 0, *t*) = 0. Following Stephens et al. ^56^, we performed principal component analysis by calculating the covariance matrix of mean-subtracted angles *θ*_*m*_(*s, t*) = *θ* (*s, t*)− ⟨*θ* (*s, t*)⟩_*t*_, defined as *θ*_Cov_(*s, s*′) = ⟨(*θ*_*m*_(*s, t*) − ⟨*θ*_*m*_⟩)(*θ*_*m*_(*s*′, *t*) − ⟨*θ*_*m*_⟩)⟩, where ⟨*θ*_*m*_⟩ is the spatial average of *θ*_*m*_(*s, t*) at a given time *t*. The eigenvalues *λ*_*n*_ and the corresponding eigenvectors *M*_*n*_(*s*) of the covariance matrix are given by Σ_*s*′_ *θ*_Cov_(*s, s*′)*M*_*n*_(*s*′) = *λ*_*n*_(*t*)*M*_*n*_(*s*). We show that superposition of four eigenvectors corresponding to four largest eigenvalues can describe the flagellum’s shape with high accuracy (see Fig. 7C)

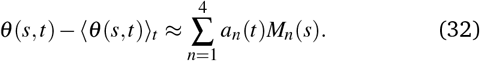

Here the four variables *a*_1_(*t*), …, *a*_4_(*t*) are the amplitudes of motion along different principal components and are given by *a*_*n*_(*t*) = ∑_*s*_ *M*_*n*_(*s*)*θ*_*m*_(*s, t*). The fractional variance of flagellum’s shape captured by *n* eigenvectors is calculated as 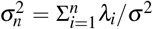 where 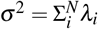. Here *N* is the total number of eigenvectors. Figure S3 shows that already two modes capture 98.6% and four modes capture 99.7% of the total variance.

## 3 Results

We fabricated axonemally-based propelled beads, by incubating isolated axonemes from *C. reinhardtii* (~10 *μ*m in length) with beads of diameter 1 *μ*m, as described in Materials and Methods (see subsection 2.1). Out of *N* =195 bead-axoneme attachments, a single bead attaches to the basal end of the axonemes in 60.5% of the cases. Attachment of beads to the distal tip occurs less frequently in 22.6% of cases, and in even less frequent events (16.9%) we observed more than one bead attached either to the tips or along the axonemal length. The attachment of the beads to the basal or the distal region of the axonemes is either symmetric, i.e. the tangent vector of the axoneme at the bead-axoneme contact point passes through the bead center, or asymmetric. We emphasize that a limitation of our 2D phase-contrast microscopy is that it is not possible to precisely distinguish a symmetric bead attachment from an asymmetric one and 3D microscopy techniques ^51^ are required to fully determine the bead-axoneme attachment geometry. This symmetric versus asymmetric attachment has consequences on the swimming dynamics, as we will discuss in the Sec. 3.3.

The propulsion of axonemally-driven beads is highly sensitive to the axonemal waveform. In our experiments, we use calcium ions at different concentrations to change the axonemal waveform and study the propulsion dynamics. In the following subsection, we summarize our results without and with calcium ions and show that calcium ions play a crucial role to achieve a directed cargo transport.

### 3.1 Overview of the results: two modes of propulsion

Returning to Fig. 1, we recall the existence of two regimes of propagation. While in the first regime, the axonemal-driven bead is propelled on a curved trajectory, in the second regime, the bead is propelled (on average) along a straight path. These two distinct modes of propulsion correspond to two different concentrations of calcium ions in the reactivation buffer (0 mM versus 1 mM), which directly affects the axonemal waveform^34,59^. As shown in Fig. 1B, in the absence of calcium ions, the time-projection of the ‘asymmetric’ axonemal shapes indicates a counter-clockwise rotation of the axoneme causing a circular swimming trajectory of the bead. However, in the presence of calcium ions, we observe a significant change in the flagellar waveform (Fig. 1C-D) that appears to be ‘symmetric’ and results in a directed bead transport.

Figure 4 characterizes in more detail the flagellar waveform of the axonemes in the two regimes observed in Fig. 1. Figure 4A-C shows a representative example of the first regime, where we have tracked the axoneme over time and quantified the curvature waves that propagate along the axonemal length. These bending waves start from the basal end (at *s* = 0), which is attached to the bead, and propagate at the frequency of *f*_0_ ~ 38.21*±*0.25 Hz toward the distal tip. Note that the beat frequency of axonemes depends on the ATP concentration, following a Michaelis-Menten kinetics with a linear trend at low amount of ATP and saturation at higher ATP concentrations around 1 mM (see Fig. S4) ^38,42^. An interesting property of these curvature waves is that if we average the flagellar shapes over one beat cycle, we obtain a circular arc with the mean curvature of *κ*_0_ around −0.19 *μ*m^−1^ (see the cyan filament in Fig. 4C). The negative sign of *κ*_0_ indicates a clockwise bend when moving from the basal end at *s* = 0 toward the distal tip at *s* = *L*. In other words, we can think of the axonemal waveform as a superposition of a traveling wave component which propagates along this circular arc, causing a counter-clockwise rotation of the axoneme and the cargo (Fig. 1A-B). However, as shown in Fig. 5A, the mean curvature of axonemes *κ*_0_ is highly sensitive to the concentration of calcium ions and almost vanishes above a calcium concentration of 0.1 mM. This mean axonemal curvature appears to be linearly related to the curvature of the swimming path of the bead, as shown by Fig. 5B. The proportionality factor is estimated to be around 2^38,60^.

**Fig. 4.**
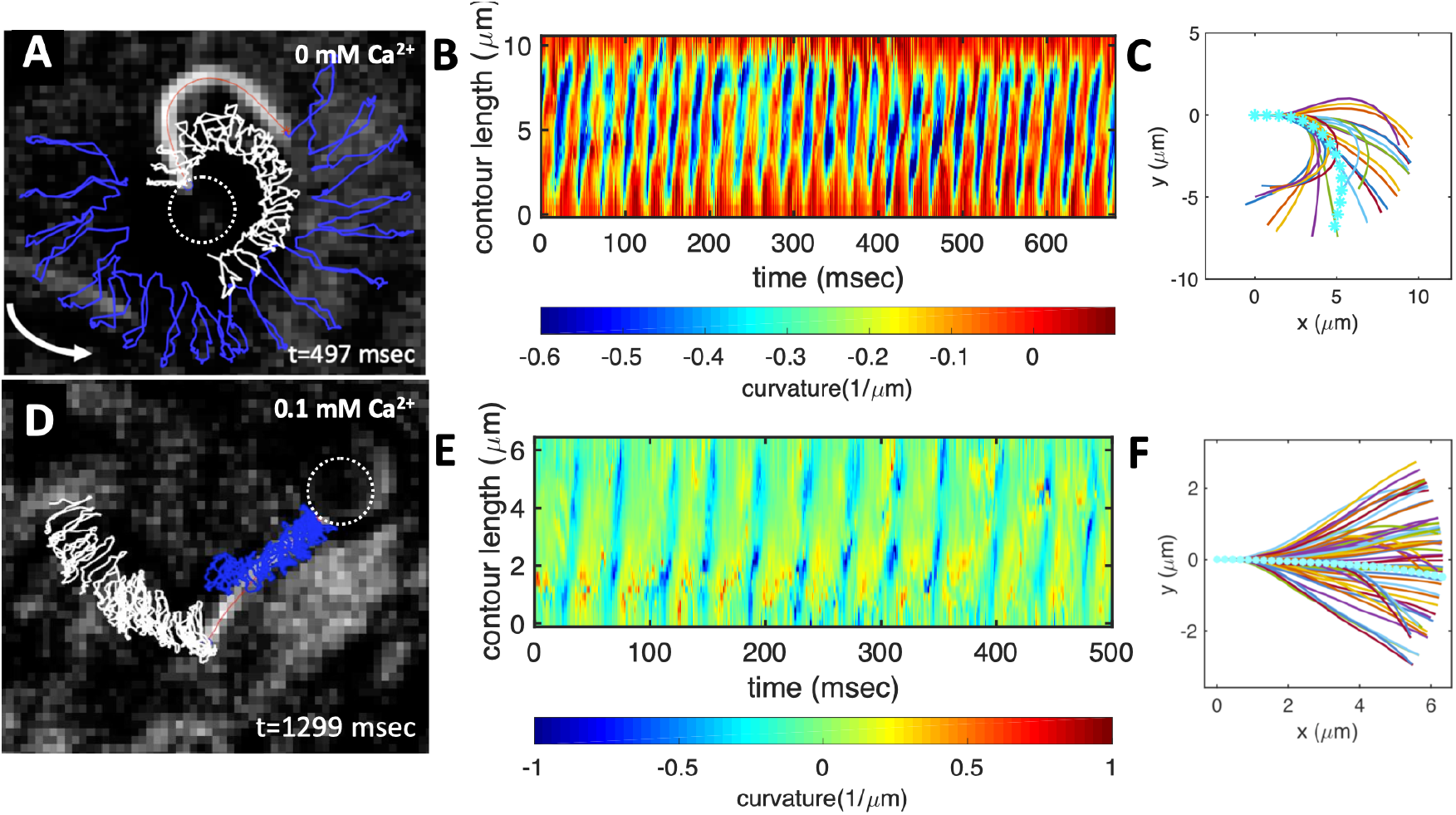
Experiments to show the effect of calcium on axonemal waveform. A) An axonemally-driven bead reactivated at [ATP]=80 *μ*M and [Ca^2+^]=0 mM (see Video 3). Tracked axoneme and trajectories of its distal (blue lines) and basal (white lines) ends. Axoneme-bead swimmer rotates counter-clockwise while the bead center follows a spiral-like path (Fig. 6B). B) Curvature waves as they travel at the frequency of 38.21 *±* 0.25 Hz from the proximal region toward the distal tip. C) Mean shape of the axoneme in cyan color averaged over one beat cycle showing a circular arc with static curvature of ~ − 0.19 *μ*m^−1^. At different time points, the axoneme is translated and rotated such that the basal end is at (0, 0) and the tangent vector at *s* = 0, which defines the **X**-axis in the swimmer-fixed frame, is along the ***x***-axis of the laboratory-fixed frame (see Fig. 1A). D) An axonemally-driven bead reactivated at [ATP]=80 *μ*M. The reactivation buffer is supplemented with 0.1 mM calcium to reduce the static curvature (see Video 4). The axoneme beats at 24.42*±*0.18 Hz and the bead-axoneme attachment appears to be symmetric. F) The axonemal mean shape with static curvature of *κ*_0_ ~-0.03 *μ*m^−1^ (filament with cyan color). The mean curvature of axoneme has dropped around ten times in comparison to axonemal shapes in panel C. This reduction in *C*_0_ causes a transition from a circular to a straight swimming trajectory.

**Fig. 5.**
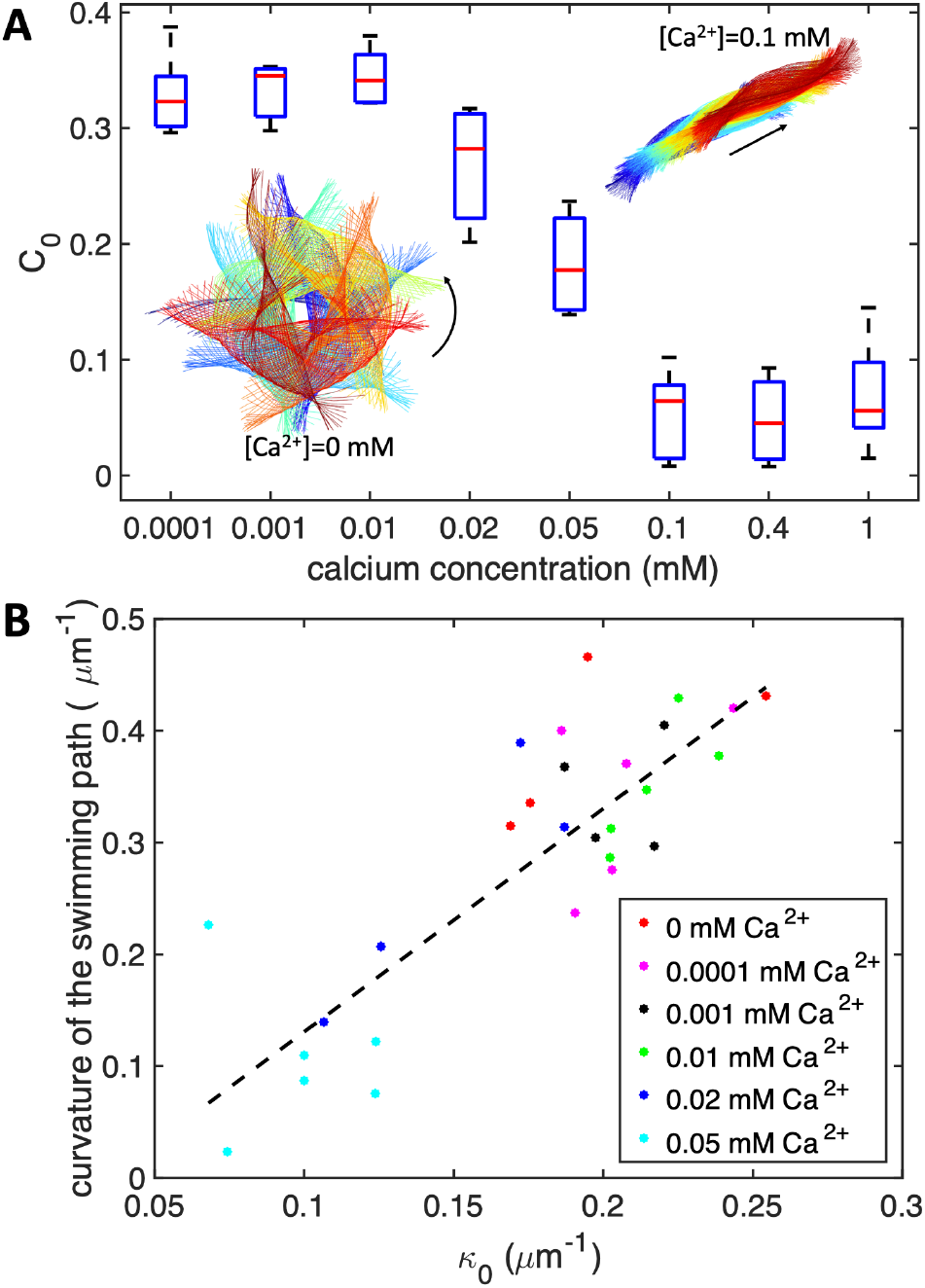
A) The dimensionless mean curvature of axonemes defined as *C*_0_ = *κ*_0_*L/*(2*π*) drops almost ten times at [Ca^2+^] around 0.01 mM. This reduction in *C*_0_ causes a transition from circular to straight swimming trajectory. The time-projection of experimental shapes of two exemplary axonemes at 0 mM and 0.1 mM [Ca^2+^] are also shown to illustrate the transition in the swimming path. At each calcium concentration, *N* = 5 axonemes (without any bead attached) are tracked to obtain *C*_0_ and the corresponding median and standard deviation. B) The curvature of the swimming path of the axonemes in panel A, plotted versus the mean curvature *κ*_0_. The slope of the linear least-square fit is close to 2.

Figure 4D-F illustrates how the axonemes beat at higher calcium concentration (0.1 mM). In this case, the axonemal waveform is composed of a propagating wave that travels along an almost straight filament, thereby propelling the axoneme and the cargo along a straight path. This reduction in *κ*_0_ decreases the rotational velocity of the axoneme compared to the experiment presented in Fig. 4A in the absence of calcium ions. As we will show in our analytical analysis, see Section 3.3, up to the leading order, ⟨Ω_*z*_⟩ scales linearly with the dimensionless mean curvature *C*_0_ = *κ*_0_*L/*2*π*. At such low values of the mean curvature of the axoneme, we achieve directional cargo transport with a mean velocity of about 20 *μ*m/sec (Fig. 1C), which is slightly slower than the migration speed of the human sperm in mucus (typically 25-45 *μ*m/sec) ^61^.

Furthermore, in most of our experiments without calcium, the axonemes rotate counter-clockwise in the microscope’s field of view with an effective 2D beat pattern, facilitating the tracking procedure. This motion is characterized by a circular swimming trajectory and a translation in the laboratory frame of reference. The combination of rotation and translation results in a spiral-like swimming path of the axoneme and the bead (see Fig. 6). Note that for the exemplary experiment in Fig. 4A, the *x*−*y* positions of the bead center exhibit tiny oscillations at the fast beat frequency of *f*_0_ ~ 38.21 Hz and a secondary slow global rotational frequency around 1.5 Hz (see Fig. 6B). As the axoneme undergoes planar shape deformations over time, at any instant of time, it may be considered as a solid body with translational and rotational velocities *U*_*x*_, *U*_*y*_ and Ω_*z*_ which we measure in the swimmer-fixed frame (Fig. S5) ^60^. These velocities oscillate in time, reflecting the fundamental beat frequency of the axoneme shown in Fig. 4A (~38.21 Hz) and its higher harmonics. In contrast, in our experiments with calcium, axonemal movement often occurs in 3D (see Video 2), making tracking of axoneme nearly impossible.

**Fig. 6.**
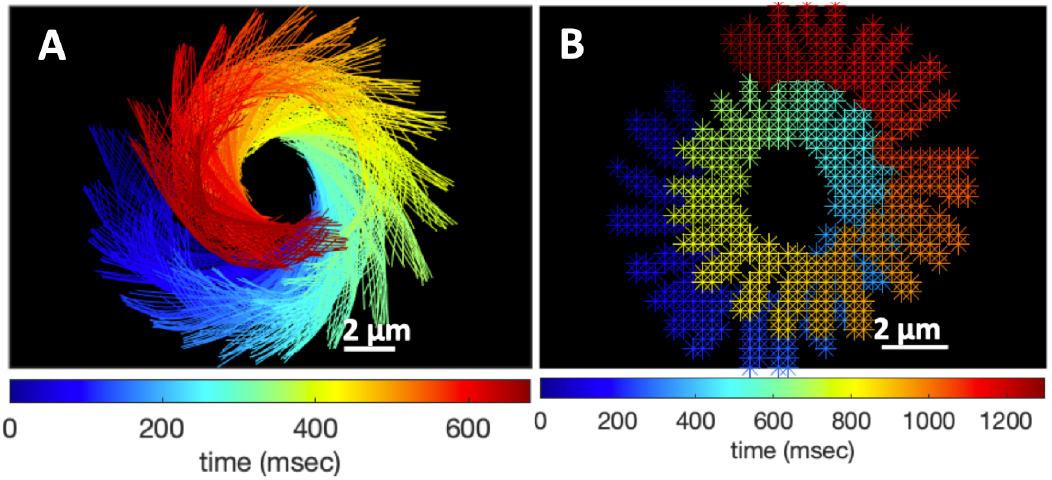
A) Color-coded time projection of the axonemal shapes of the axoneme presented in Fig. 4A ([Ca^2+^]=0 mM). B) Color-coded trajectory of the bead center from Fig. 4A showing a spiral-like trajectory. The global rotational frequency of the swimmer is around 1.5 Hz (~ 1 full rotation in around 650 msec; see Videos 3,5)

Finally, we also performed experiments with larger diameter beads to explore how bead size affects beating frequency and mean curvature of the axonemes. A summary of the beat frequency and intrinsic curvature of axonemes (all reactivated at [ATP]=200 *μ*M and [Ca^2+^]=0 mM) propelling beads of diameter 2 and 3 micron is given in supplemental Table S1. In these experiments, as we increased the bead diameter from 1 *μ*m to 3 *μ*m, we observed a reduction at both the averaged beat frequency (~ 12%) and the intrinsic curvature (~ 31%) of the axonemes.

### 3.2 Description of the flagellar shapes

Having described the two modes of propulsion with and without calcium, we now focus on the principal component analysis (PCA) of the flagellar waveform as described in subsection 2.4^39,56^. In our flagella-driven beads, the propulsive force and torque exerted on the bead depends on the axonemal waveform. Using the *x* and *y* coordinates of the tracked flagellum (see subsection 2.2), we calculate *θ* (*s, t*) which is defined as the angle between the local tangent vector of the centerline of the tracked flagellum and the ***x***-axis (see Fig. 1A). We then average the axonemal shapes over one beat cycle to obtain the mean shape of the axoneme, which corresponds to the arc-shaped filament colored in cyan in Fig. 4C.

Figure 7A shows the color map of the covariance matrix of the axoneme shown in Fig. 4A, which has an effective reduced dimensionality with a small number of non-zero eigenvalues. We note that four eigenmodes *M*_*n*_(*s*) (*n* = 1,.., 4) corresponding to the first four largest eigenvalues of *θ*_Cov_(*s, s*′) capture the axonemal shapes with very high accuracy (see Video 6). These four eigenvectors are plotted in Fig. 7B. We reconstruct the axonemal shapes using the four dominant modes and the corresponding time-dependent motion amplitudes *a*_*n*_(*t*) (*n* = 1,.., 4) using Eq. 32. The reconstructed shapes are presented in Fig. 7C, where the green dashed lines show the experimental data. The contribution of different modes in reconstructing the axonemal shapes is shown in Fig. S6. Please note that higher spatial modes in combination with the corresponding time-dependent motion amplitudes, i.e. *a*_2_(*t*)*M*_2_(*s*), *a*_3_(*t*)*M*_3_(*s*) and *a*_4_(*t*)*M*_4_(*s*), are approximately standing waves. The motion amplitudes *a*_1_(*t*) and *a*_2_(*t*), which oscillate at the axonemal beat frequency *f*_0_, are shown in Fig. 7D; see also Fig. S7 for a plot of motion amplitudes *a*_1_(*t*) to *a*_4_(*t*) versus each other.

**Fig. 7.**
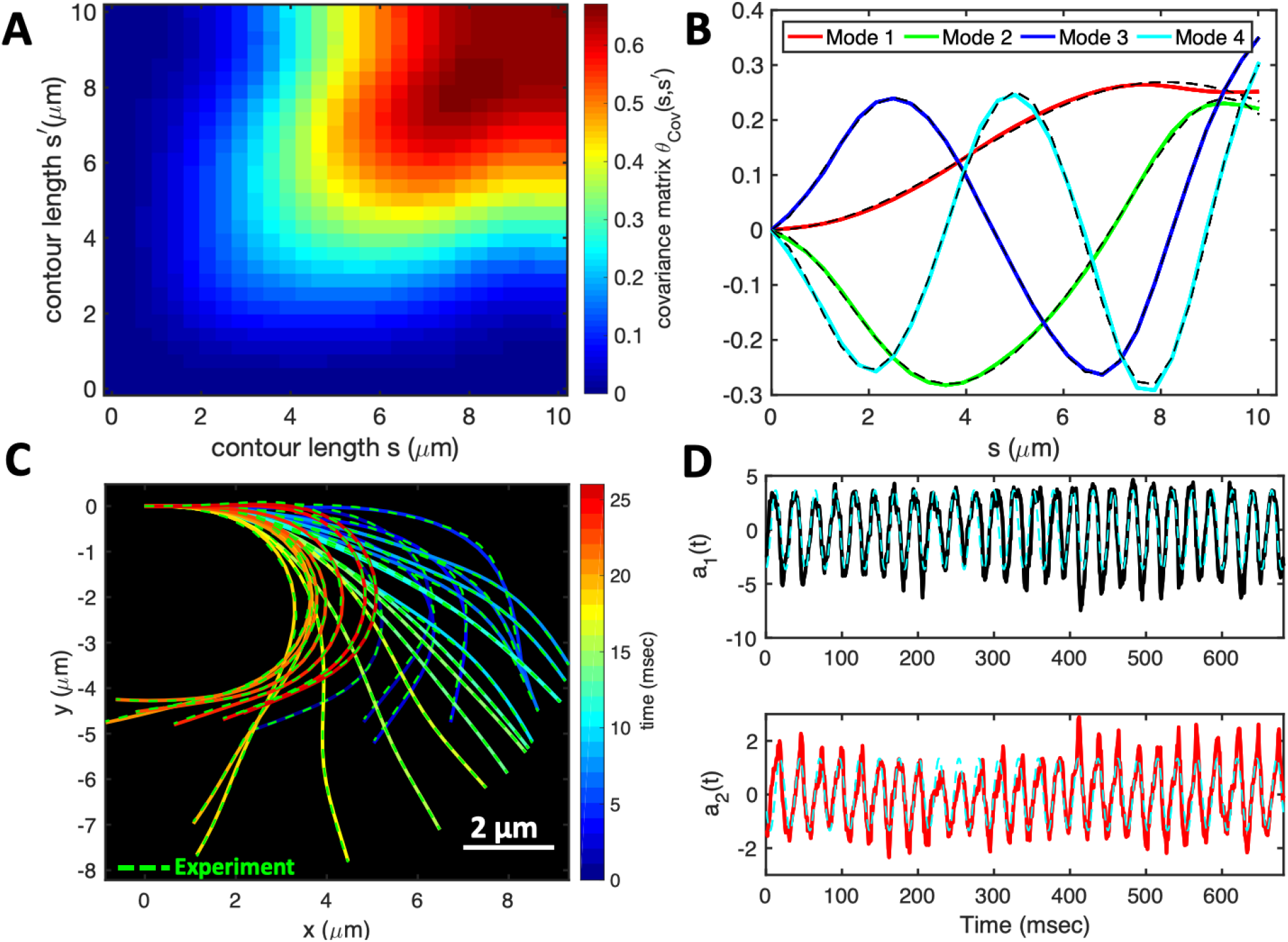
PCA analysis of the axoneme presented in Fig. 4A. A) The covariance matrix *θ*_Cov_(*s, s*′) of fluctuations in angle *θ* (*s, t*). B) The four eigenvectors corresponding to the first four largest eigenvalues of *θ*_Cov_(*s, s*′). C) The time-projection of the experimental data and the corresponding shapes reconstructed using the four eigenmodes presented in panel B. D) The time-dependent motion amplitudes *a*_1_(*t*) and *a*_2_(*t*) as defined in Sec. 2.4. The black and cyan dashed lines in panel B and D show the Fourier fits.

### 3.3 Analytical approximations of rotational and translational velocities of an axonemal-propelled bead

In this section, we discuss analytical approximations for the mean translational and rotational velocities of a flagellar-propelled bead in the regime of small wave amplitude. To describe the traveling curvature waves analytically, we use the following simplified waveform which is composed of a traveling wave component with amplitude *C*_1_ superimposed on a circular arc with the mean curvature of *C*_0_

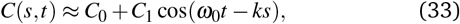

where *ω*_0_ = 2*π f*_0_, *k* = 2*π/λ* is the wave number. For our exemplary axoneme in Fig. 4A, following the method described in Ref. ^38^, we calculate the wavelength to be *λ* ~13.19 *μ*m (see Fig. S8), which is ~ 27% larger than the axonemal contour length *L* ~ 10.38 *μ*m.

Next, we obtain analytical expressions for the mean translational and rotational velocities of the swimmer in the regime of small *C*_0_ and *C*_1_. For our analytical approximations and simulations, we used the resistive-force theory (RFT) which is presented in Sec. 2.3. To check the validity of RFT, we extracted rotational and translational velocities of an exemplary axonemal-driven bead shown in the supplemental Video 7 and compared the results with velocities obtained by RFT simulations using experimental beat patterns (see Sec. 2.3). This comparison is presented in Fig. S9, which demonstrates a semi-quantitative agreement between RFT simulations and experimental data (see also Sec. 3.4). In the rest of the section, we will use the simplified waveform, as defined in Eq. 33, to perform analytical analysis as well as numerical simulations for two cases: first, for a symmetric bead-axoneme attachment, and second, for an asymmetric attachment.

#### 3.3.1 Symmetric bead-axoneme attachment

Let us first consider the example where an axoneme is attached symmetrically from the basal side to a bead, so that the tangent vector at *s* = 0 passes through the bead center. To determine the dependence of the translational and rotational velocities of the swimmer on the dimensionless bead radius *r* = *R/L*, we used the prescribed form of the curvature waves in Eq. 33 and imposed the force-free and torque-free conditions in 2D. The drag matrices of bead and axoneme are calculated with respect to the reference point, which is defined as the bead-axoneme contact point (see Sec. 2.3.1). We approximate analytically the averaged angular and linear velocities in the swimmer-fixed frame, as defined in Fig. 1A, in the limit where the wavelength of the beat pattern, *λ*, is equal to the size of the axoneme, *L*, and up to the first order in *C*_0_, and the second order in *C*_1_ to obtain

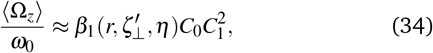

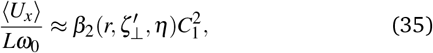

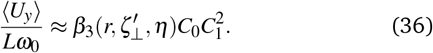

The full expressions of functions *β*_1_, *β*_2_ and *β*_3_ are presented in the supplemental information, Eqs. S.1-S.3. The results for a bead of size *r* = 0.1 with drag anisotropy *η* ≡ *ζ*_‖_ */ζ*_⊥_ = 0.5 simplifies to

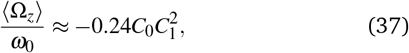

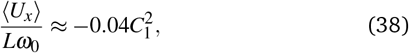

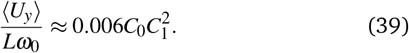

We note that in Eqs. 34-36, without the even harmonic, i.e intrinsic curvature *C*_0_ = 0, the axoneme swims in a straight path with ⟨*U*_*y*_⟩ = 0, ⟨*U*_*x*_⟩ proportional to the square of traveling wave component *C*_1_^60,62,63^, and the mean rotational velocity ⟨Ω_*z*_⟩ vanishes (see the solid lines in Fig. 8A-F and Fig. S10).

**Fig. 8.**
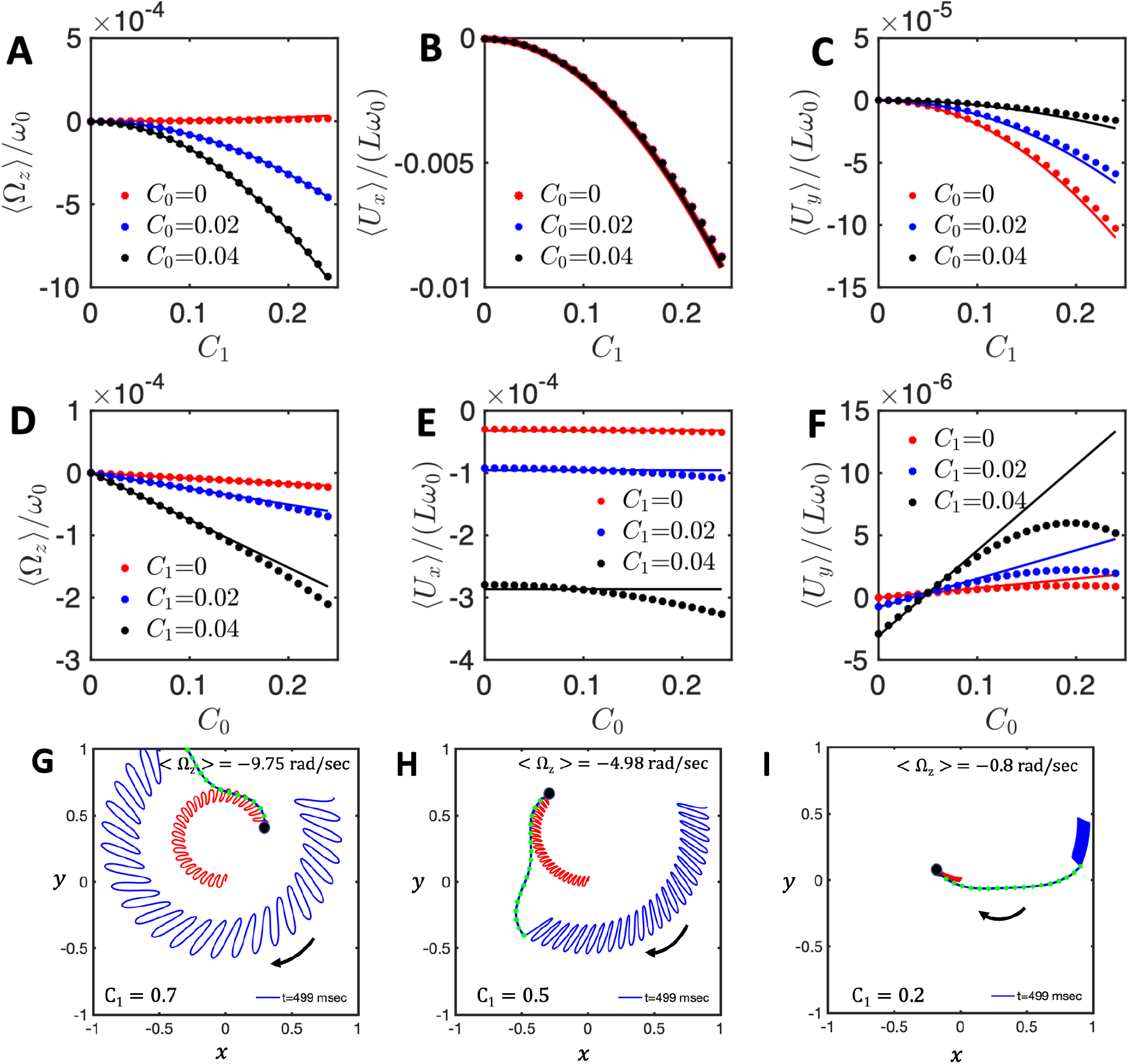
A-F) Comparison of analytical approximations of rotational and translational velocities with numerical simulations at bead radius of *R* = 0. Solid lines present the analytical approximations summarized in Eqs. 34-36 and dots show the numerical simulations. G-I) Numerical simulations performed with the simplified waveform to show the effect of *C*_1_. A bead of radius *R/L* = 0.1 is attached symmetrically to a model flagellum. The mean rotational velocity decreases as the amplitude of dynamic mode *C*_1_ decreases from G) *C*_1_ = 0.7, to H) *C*_1_ = 0.5 and further to I) *C*_1_ = 0.2. The static curvature is fixed at *C*_0_ = 0.2.

In parallel to our analytical approximations, we also performed numerical simulations given the simplified waveform in Eq. 33 and using RFT with the realistic value of *η* = *ζ*_‖_ *ζ*_⊥_ = 0.5. As shown in Fig. 8A-F, the comparison between numerical simulations and the full analytical approximations presented in Eqs. S.1–S.3, shows good agreement at small values of *C*_0_ and *C*_1_ and deviations at larger values. In addition, three exemplary simulations in Fig. 8G-I highlight the dependency of the mean rotational velocity ⟨Ω_*z*_⟩ of the model swimmer on the square of the traveling wave component *C*_1_.

To investigate the dependency of the mean translational and rotational velocities of our model swimmer on the bead size, we plotted Eqs. 34-36 as a function of *r* (Fig. 9). We also performed numerical simulations at different bead radii. As shown in Fig. 9, there is a good agreement between our numerical simulations and analytical approximations at small values of *C*_1_ (panels A-C) but deviations at larger values (panels D-F). Furthermore, it is remarkable that while ⟨*U*_*x*_⟩ reduces monotonically with the bead radius *r*, both ⟨Ω_*z*_⟩ and ⟨*U*_*y*_⟩ exhibit a non-monotonic dependence. This anomalous behavior is counter-intuitive, since rotational and translational speeds are expected to decrease with increasing the bead radius. To examine this anomalous behavior, we looked at the asymptotic expressions of Eqs. 34-36 in two opposite limits of small and large bead radii. For the case where the bead radius *R* is much larger than the flagellar length *L* (small 1*/r*), we obtain up to the second order in *r*^−1^

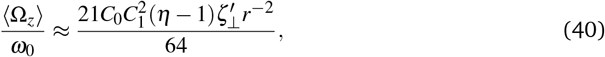

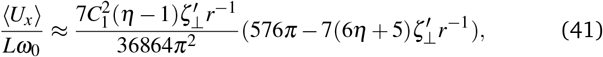

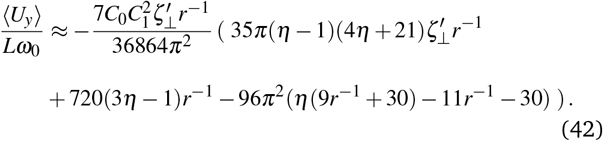

**Fig. 9.**
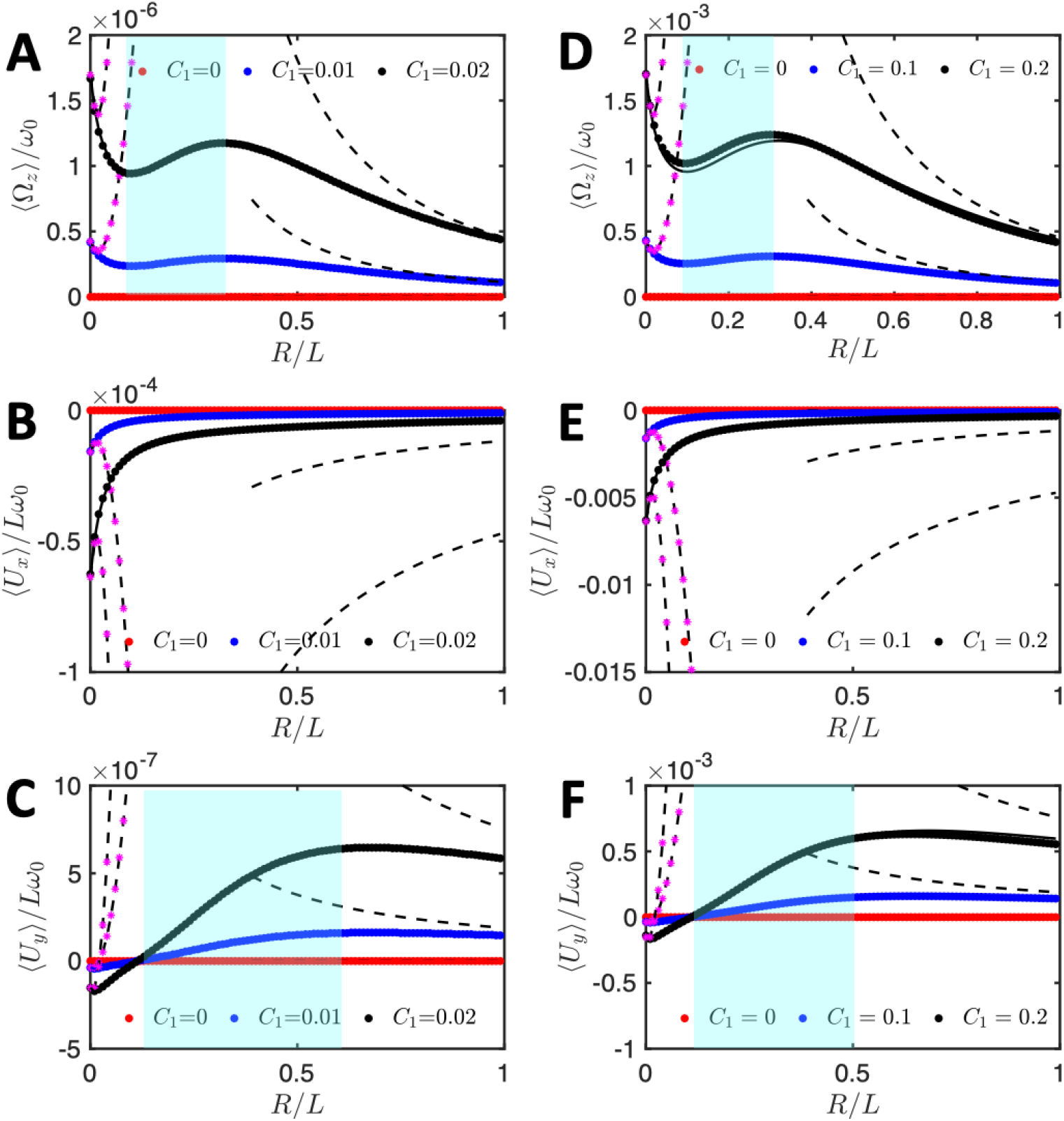
Anomalous flagella-based propulsion speed of a symmetrically attached bead as a function of its dimensionless radius *R/L*. Contrary to expectations, in the cyan region the mean translational and rotational velocities increase with increasing the bead radius. Analytical approximations (continuous lines calculated from Eqs. S.1-S.3) and simulations (dotted points) are performed at different values of *C*_1_, while the intrinsic curvature of the axoneme is fixed at *C*_0_ = −0.01 in (A-C), and *C*_0_ = −0.1 in (D-F). The black dashed curves show the trend expected in the limit of large bead radius (*r* = *R/L >* 1), as presented in Eqs. 40-42. The black dashed lines with stars in magenta illustrate the opposite limit of the small *r*, as given in Eqs. 43-45.

In the opposite limit of small *r* (i.e. *R ≪ L*), up to the second order in *r*, we set *η* = 0.5 and 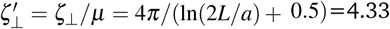 to get

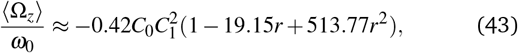

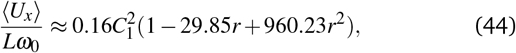

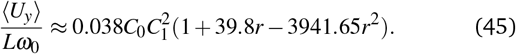

The black dashed lines (with and without stars in pink color) in Fig. 9 show these asymptotic results. The transition from *r*^2^ dependency at small values of bead radius to *r*^−2^-trend at large values of *r* manifests itself in an anomalous behavior. Finally, we note that in the absence of the bead (*r* = 0), Eqs. 43-45 reduce to Eqs. S.4-S.6 as discussed in the Refs. ^39,40^.

#### 3.3.2 The sideways bead-axoneme attachment contributes to the rotational velocity of the swimmer

In our experiments, we frequently observed an asymmetric bead-axoneme attachment, i.e. the tangent vector of the axoneme at *s* = 0 does not pass through the bead center. This case is schematically shown in Fig. 2A and Fig. 10B. Care needs to be taken since, as mentioned previously, a bead that appears to be symmetrically attached to an axoneme in our 2D phase-contrast images, could in reality be attached asymmetrically and 3D imaging techniques are necessary to distinguish these two scenarios. Interestingly, our analytical approximations and simulations confirm that this asymmetric bead-axoneme attachment is enough to rotate the axoneme and presence of the static curvature or the second harmonic are not necessary for rotation to occur. For this analysis, we consider the 2D geometry where the center of the bead is at position *X*_*B*_ and *Y*_*B*_ which is measured with respect to the coordinate system defined at the bead-axoneme contact point (Figs. 2A and 1A). The drag matrix of the bead is given by Eq. 15. To focus on the effect of the sideways bead attachment on the axonemal rotational velocity, we set the intrinsic curvature *C*_0_ to zero and consider a model sperm-like axoneme with only the traveling wave components *C*_1_ ≠ 0 in Eq. 33. We calculate the mean rotational velocity of the axonemal-driven bead by combining the drag matrix of the bead and the axoneme (see Materials and Methods) to arrive at

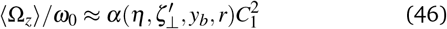

where dimensionless quantities are *r* = *R/L, y*_*b*_ = *Y*_*B*_*/L, η* = *ζ*_‖_ */ζ*_⊥_ and 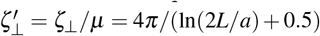. Here *μ* is the dynamic viscosity of the ambient water-like fluid and *a* = 100 nm is the axonemal radius. Note that the variable 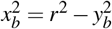 is not an independent variable. For the complete form of the function *α*, see Eq. S.7.

**Fig. 10.**
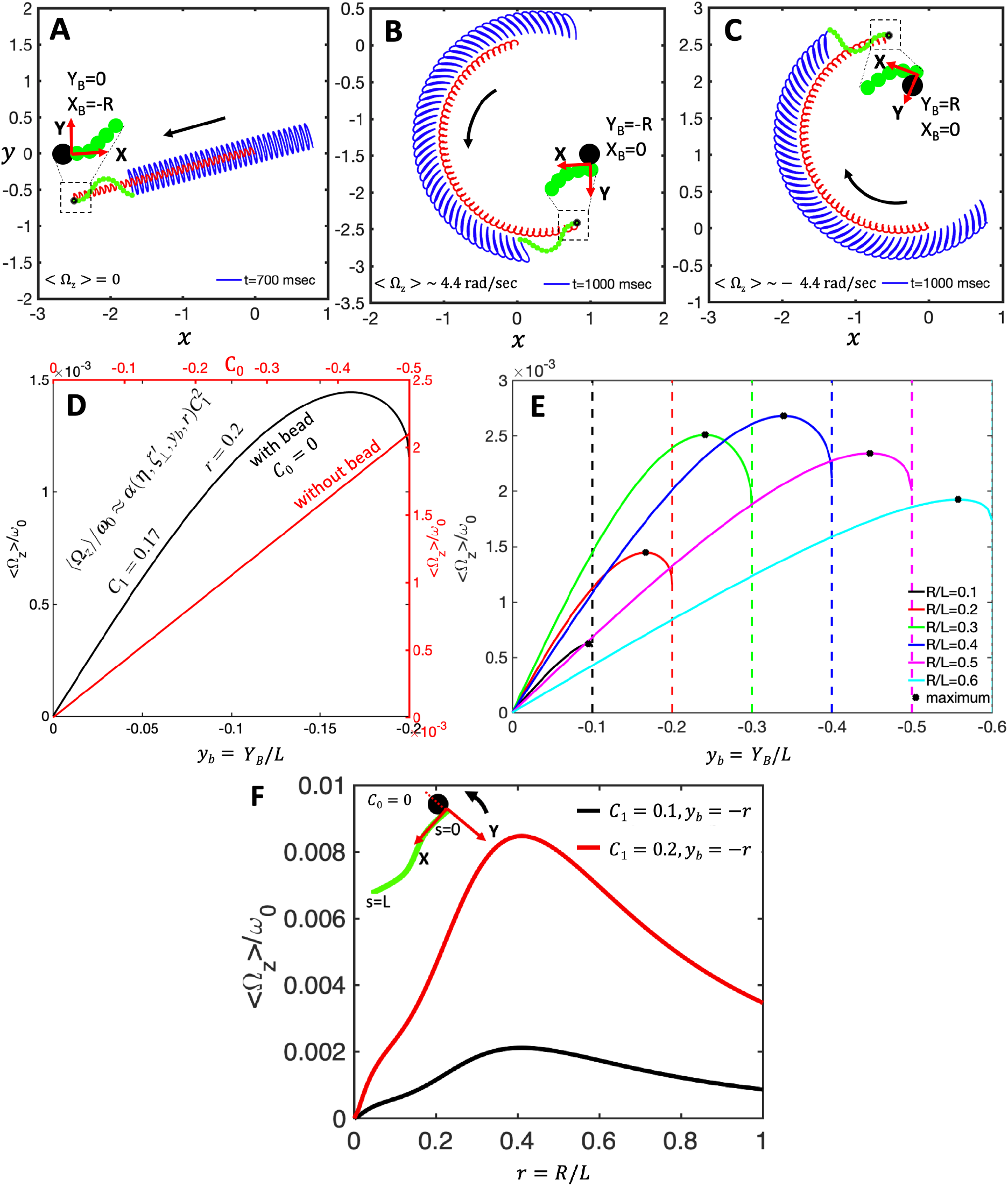
A-C) Asymmetric versus symmetric bead attachment to an axoneme with a beat pattern consisting only of the traveling wave component *C*_1_. While the axoneme in panel A, to which the bead is symmetrically attached, swims on a straight path (Video 8), the axonemes in panel B (Video 9) and C (Video 10), with asymmetric bead attachment, swim on curved paths. D) Comparison of the effect of the asymmetric bead attachment (in black) as a function of *y*_*b*_ versus the effect of the intrinsic curvature *C*_0_ (in red) on ⟨Ω_*z*_⟩. E) The averaged angular velocity ⟨Ω_*z*_⟩ changes non-monotonically with *y*_*b*_ for different bead radii. *X*_*B*_ and *Y*_*B*_ are the coordinates of the bead center in the swimmer-fixed reference frame. Parameters are *η*=0.5, 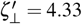 and *C*_1_ = 0.1. F) The mean rotational velocity of a bead which is asymmetrically attached to an axoneme with *y*_*b*_ = −*r* and *x*_*b*_ = 0 for two different values of *C*_1_. The mean curvature *C*_0_ is set to zero.

Some important consequences of Eq. 46 are as follows: First, since the intrinsic curvature *C*_0_ is set to zero, ⟨Ω_*z*_⟩ is zero for the case of a symmetric bead attachment, i.e. *x*_*b*_ = −*R/L* and *y*_*b*_ = 0. The effect of sideways bead attachment on rotational velocity of the axoneme is highlighted in Fig. 10A-C. In these simulations, the model axoneme has only the traveling wave component *C*_1_ and it swims in a straight path if the bead is attached symmetrically (Fig. 10A and Video 8). An asymmetric bead-axoneme attachment causes the axoneme to rotate (Fig. 10B-C and Videos 9-10). Second, a comparison between the effect of the intrinsic curvature (Eq. S.1 with *r* = 0) versus asymmetric bead attachment (Eq. 46 with *η* = 0.5, 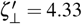 and *r* = 0.2) on rotational velocity shows that the contribution of the asymmetry in the attachment to ⟨Ω_*z*_⟩ is comparable to that of the intrinsic curvature, *C*_0_, see Fig. 10D. Third, as shown in Fig. 10E, the maximum rotational velocity occurs in the vicinity of *y*_*b*_ = −*r*. Last, ⟨Ω_*z*_⟩ changes non-monotonically with the bead radius as shown in Fig. 10F. Note that for an asymmetric attachment of the bead at *y*_*b*_ = −*r*, the maximum ⟨Ω_*z*_⟩ occurs around *r* = 0.4.

### 3.4 Analysis with the experimental waveform

To confirm that anomalous propulsion regime and the rotation induced by asymmetric cargo attachment is general and not limited to the simplified waveform introduced in Eq. 33, we also used the experimental beat patterns of the exemplary axoneme in Fig. 4A. We performed RFT simulations to compute mean translational and rotational velocities of an axonemal-propelled bead using the experimental waveform as an input for both asymmetric and symmetric bead-axoneme attachment. As shown in Fig. 11, the anomalous propulsion regime exists for both symmetric and asymmetric attachments. Depending on the sign of the static curvature of the axoneme *C*_0_ and the position at which the bead is attached, the asymmetric attachment may contribute to an increase or decrease in the overall mean rotational velocities. For the axoneme in Fig. 11, *C*_0_ is negative and the sideways bead attachment at *Y*_*B*_ = *R* (Fig. 11A) acts against the rotation induced by the intrinsic curvature *C*_0_. The opposite happens in Fig. 11B where the bead attachment at *Y*_*B*_ = −*R* amplifies the rotational velocity of the axoneme. We also note that the anomalous propulsion regime is more pronounced in panel B where the bead is attached sideways at *Y*_*B*_ = −*R*. Note that for the symmetric bead-axoneme attachment, the general trend shown in Fig. 11C is consistent with our analytical approximations presented in Fig. 9.

**Fig. 11.**
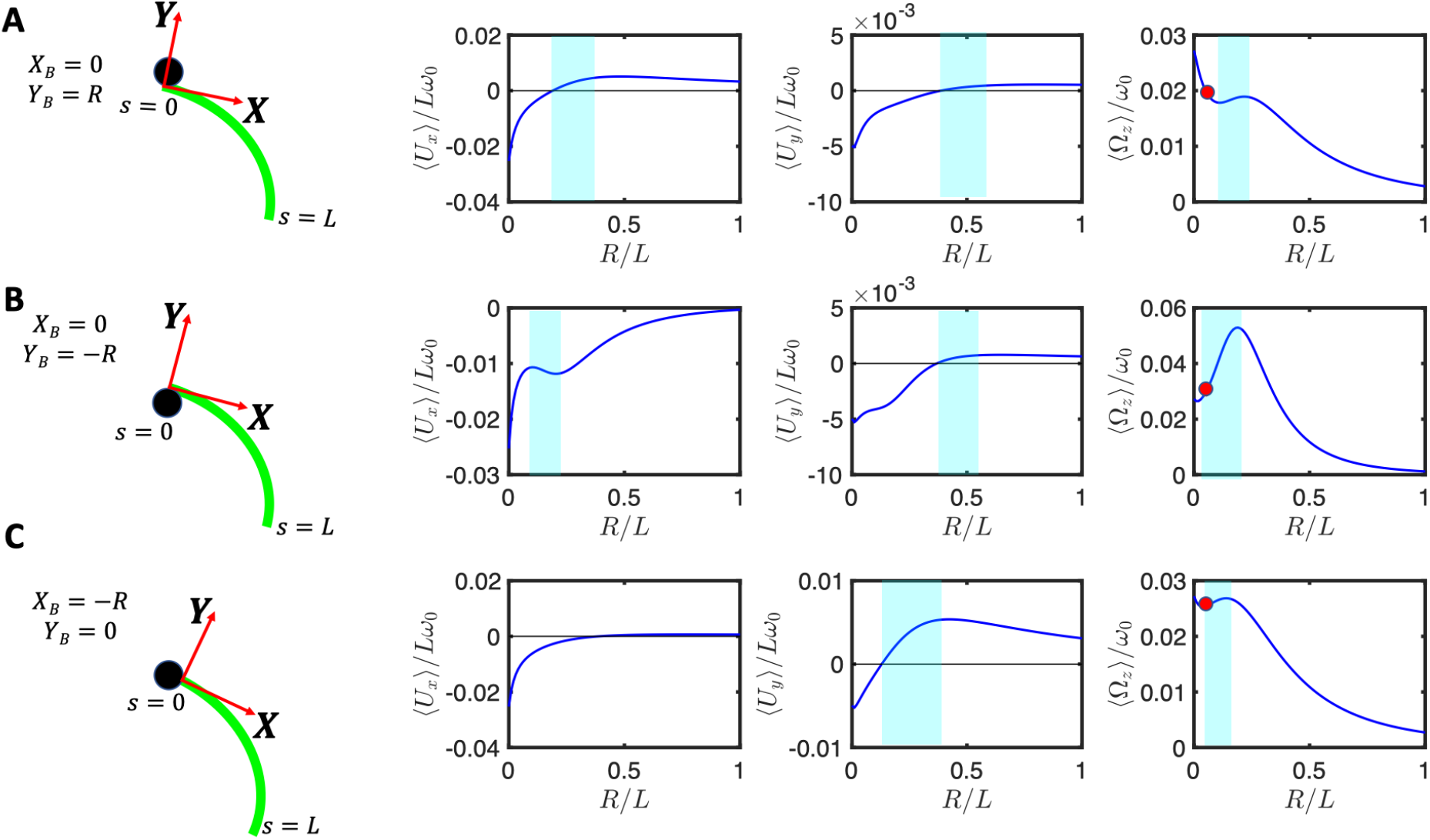
Experimental beat pattern in Fig. 4A are used to calculate translational and rotational velocities of an axonemal-propelled bead attached (A) asymmetrically at *Y*_*B*_ = *R, X*_*B*_ = 0 (B) asymmetrically at *Y*_*B*_ = −*R, X*_*B*_ = 0 and (C) symmetrically at *Y*_*B*_ = 0, *X*_*B*_ = −*R*. Anomalous propulsion regimes are highlighted in cyan color. Red circles mark the experimental bead size of *R/L* =0.05 and the corresponding rotational velocities. Note that the trend observed in panel C is consistent with the trend predicted by our analytical calculations with a simplified waveform for a symmetric bead-axoneme attachment as shown in Fig. 9.

An experimental confirmation of our theory is provided by the explicit comparison between the measured rotational velocity, ⟨Ω_*z*_⟩*/ω*_0_, with the predictions using RFT theory. To this end, we have indicated by a red circle in the right most panels of Fig. 11A-C the measured values as a function of the bead size used, *R* = 0.5 *μ*m (so the ratio *R/L* = 0.05). The value of ⟨Ω_*z*_⟩*/ω*_0_ = 0.026, reported in Fig. 11C, was documented in Fig. S5C (see the red line). In this case, the axoneme, as shown in Fig. 6A, globally rotates around 2*π* in the time interval of ~650 msec, which with *f*_0_ = 38.21 Hz, results to ⟨Ω_*z*_⟩*/ω*_0_ ~0.04. This value is closest to the value of 0.029 in panel B with an asymmetric bead-axoneme attachment. Overall, we observe a good agreement between the predicted and the experimentally measured values of ⟨Ω_*z*_⟩*/ω*_0_.

We also considered the beat pattern from our experiments at higher calcium concentration, corresponding to Fig. 4D, to study the propulsion of a model bead at different radii with symmetric versus asymmetric bead-axoneme attachment. Figure 12 also shows the existence of an anomalous propulsion regimes as highlighted by the bands in cyan color in the three graphs on the right. The experimental value of ⟨Ω_*z*_⟩*/ω*_0_ ~ 0.004 (total rotation of ~ *π*/4 in 1299 msec; see Video 4) is slightly larger than the values 0.0024, 0.0029 and 0.0028 in panels A-C, respectively, which are highlighted by the red circles in Fig. 12A-C. Finally, comparing Figs. 11 and 12 shows that, as expected, the dependence of the mean translational and rotational velocities on the bead size is highly sensitive to the flagellar waveform.

**Fig. 12.**
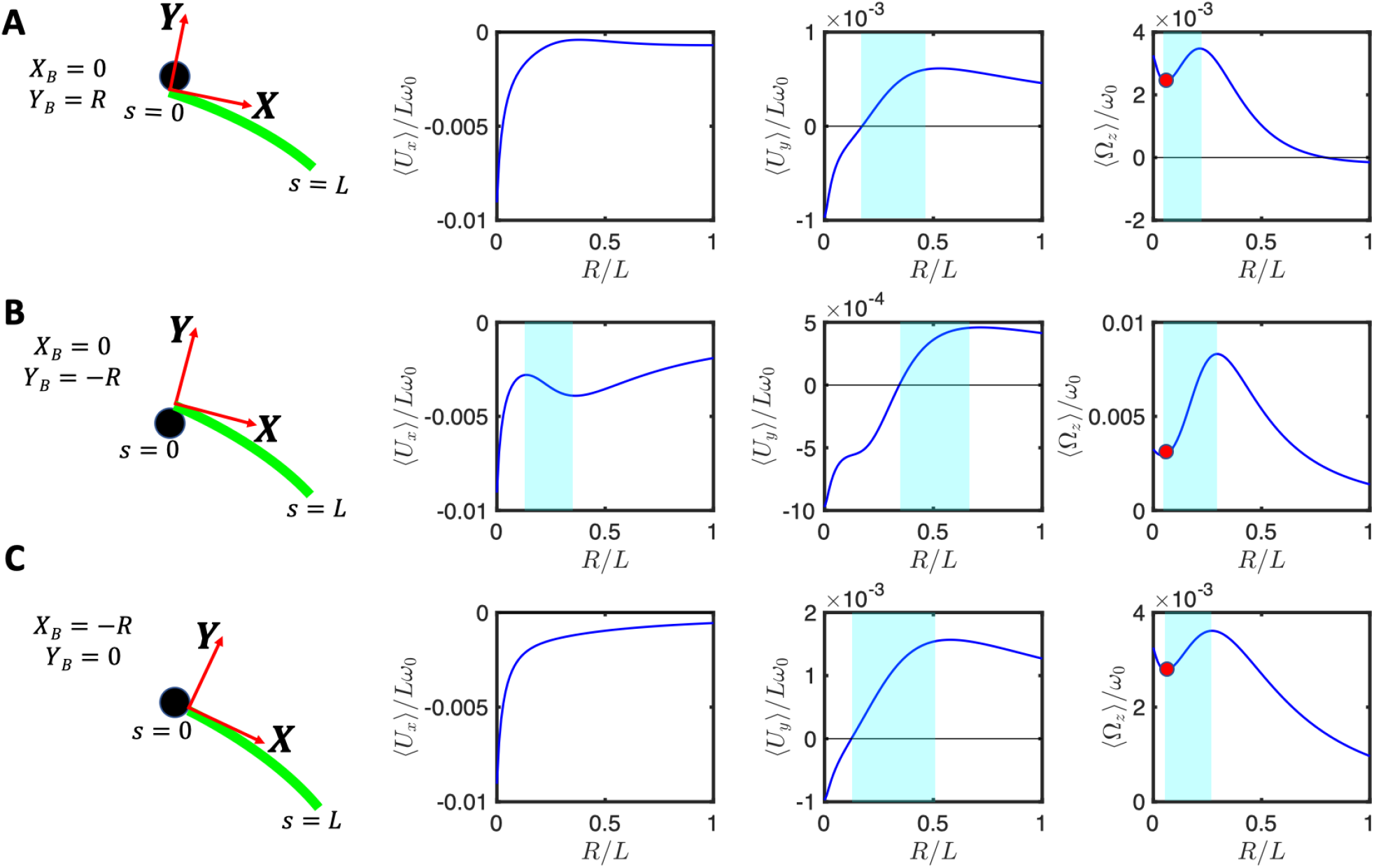
Experimental beat pattern shown in Fig. 4D with 0.1 mM [Ca^2+^] and [ATP]=80 *μ*M are used to calculate the mean translational and rotational velocities of an axonemal-driven bead attached (A) asymmetrically at *Y*_*B*_ = *R, X*_*B*_ = 0 (B) asymmetrically at *Y*_*B*_ = −*R, X*_*B*_ = 0 and (C) symmetrically at *Y*_*B*_ = 0, *X*_*B*_ = −*R*. Anomalous propulsion regimes are highlighted in cyan color. Red circles mark the experimental bead radius of *R/L*=0.05 and the corresponding values of ⟨Ω_*z*_⟩*/ω*_0_ (see supplemental Video 4). Note that the general trend in panel C is consistent with the analytical analysis presented in Fig. 9 for a symmetric bead attachment.

## Conclusions

In this work, we have characterized the motion of a bead (a cargo) propelled by an ATP-reactivated axoneme isolated from the green algae *C. reinhardtii*. We observed two distinct regimes of bead propulsion depending on the calcium concentration. The first regime describes the bead motion along a curved trajectory which is observed in experiments at zero or very small concentration of calcium ions (less than 0.02 mM). A directed bead propulsion along straight trajectories occurs at higher calcium concentrations where the cargo is propelled at an averaged velocity around 20 *μ*m/sec. This is comparable to the typical human sperm migration speed in mucus ^61^. High resolution structural information obtained by electron cryotomography ^64^, strongly supports the idea that calcium could regulate the transmission of mechanosignals. Gui et al. ^64^ have shown the presence of calmodulin, a calcium responsive protein, at the interface between RS1 (radial spoke 1) and IDAa (inner dynein arm a). Calcium may induce a conformational change in calmodulin, which may directly alter the wave pattern by affecting RS1-IDAa interaction. An alternative plausible mechanism is that calcium affects a calmodulin-like subunit (LC4) of the outer dynein arm (ODA) and consequently, influence the dynein behavior ^65^. Further experiments are required to clarify the mechanism of dynein regulation and the precise role of calcium in shaping the flagellar waveform.

Since the beads in our experiments were incubated with demembranated axonemes, the quantity, spacing and location of beads attached to the axonemes were not controlled. Normally, a small fraction of the beads (less than 10%) adheres to the axonemes at random sites preferably at the basal or the distal ends; see Videos 11-15 for examples of having two axonemes attached to one bead, two beads attached to one axoneme at two opposite ends, one bead along the contour length, one bead at the distal tip, and finally two beads at one end.

The axonemes beat with an asymmetric waveform, which resembles the forward swimming motion of flagella in intact cells ^24^. We extracted the axonemal shapes using gradient vector flow technique^49,50^ and quantitatively described the beating patterns by the dimensionless curvature *C*(*s, t*) at time *t* and at arc-length *s* along the axonemal length. Our PCA analysis shows that the axonemal motion is described with a high degree of accuracy, taking into account only the first four dominant eigenmodes corresponding to the first four largest eigenvalues. In this paper, we are focusing on the examples where axoneme-bead swimmer swims effectively in 2D in the vicinity of the substrate. This greatly facilitates the tracking of axonemes and the data analysis. However, we also observed examples where the axoneme-bead swimmer undergoes a tumbling motion in 3D, as shown in the supplemental Videos 16-18. This out of plane swimming dynamics complicates the tracking process of the axonemes. In future studies, 3D microscopy techniques^51^ are necessary to capture the full 3D swimming dynamics of the isolated axonemes.

We used a simplified waveform to describe the axonemal shapes which is composed of a traveling wave component propagating along a circular arc (Eq. 33). This simplified waveform allows us to obtain analytical expressions for translational and rotational velocities of an axonemal-propelled bead in the limit of small amplitudes of curvature waves. The rotational velocity of an axoneme is predominately controlled by its static curvature *C*_0_. Remarkably, our analysis with the simplified waveform predicts an anomalous propulsion regime where the rotational and/or some of the translational velocity components of an axonemal-propelled bead increase with the bead radius (Fig. 9). We also used the experimental beat patterns to demonstrate that this counter-intuitive regime is not limited to the simplified waveform and also exists for waveforms closer to the experimental ones. This anomalous propulsion regime has also been predicted for a model sperm-like swimmer which has a zero mean curvature, and is propelled by a traveling wave motion, as illustrated by Fig. 10F. Consistent with this, we also observed anomalous regimes in our experiments with an increased calcium concentration (0.1 mM instead of 0 mM), which result in a significant reduction (approximately 10 times) in the mean curvature of the axonemes; see Figs. 5.

An anomalous cargo transport regime was also predicted in biofilm forming bacteria *Pseudomonas aeruginosa* (PA14) ^11,12^, where swimming is driven by multiple (on average two) rotating helical flagella (length~4 *μ*m) that can bundle to propel the bacterium in a corkscrew-like motion or unbundle to change direction, exhibiting a run-and-tumble swimming pattern^66^. This anomalous behavior is expected to exist for a hypothetical mutant of PA14 which has a larger (around three times) bacteria size in comparison to the wild type. This up-scaling of bacteria size results in a larger rotational drag coefficient of flagella than bacterial body, which in Ref. ^11^ is found to be the criteria for the anomalous propulsion. Whether such a hypothetical largescale bacterium exists is an open question. However, this anomalous propulsion regime could be important in bacterial swimming in polymeric solutions, where due to steric interactions between flagella and polymers, the rotational drag coefficient of the flagella can become larger than the bacterial body, fulfilling the criteria for anomalous propulsion. Thus, as experimentally observed and contrary to our expectations, the swimming speed of bacteria in the polymeric solutions can increase^67,68^. In our system, however, we are not able to obtain a simple analytical criteria for the anomalous propulsion as it results from the full calculations which include inverting the 3 *×* 3 drag matrix of the bead-axoneme swimmer. From a general physics perspective, we remark that the anomalous regime corresponds to a change in the partitioning between translation (in the two physical directions) and rotation as *R* increases, so that some components may increase with *R* over a range while other components decrease, as dictated by the overall increase in the drag of the bead.

Finally, our analysis shows that asymmetric cargo-axoneme attachment contributes at the same order of magnitude as the mean curvature to the rotational velocity. In other words, a sperm-like beating flagellum without mean curvature and second harmonic swims in a curved trajectory if it is attached sideways to a cargo. In our experiments, since we image the sample in 2D, we are not able to precisely distinguish symmetric versus asymmetric bead-axoneme attachments. In a 2D-projected image, a symmetric bead-axoneme attachment could in reality be an asymmetric one. Moreover, as the bead-axoneme swimmer goes slightly out of focus, the attachment in some frames seems to be symmetric and in other frames asymmetric. The 3D microscopy techniques, similar to the one used in Ref. ^51^ are necessary to fully characterize symmetric versus asymmetric axoneme-bead attachment in 3D. This 3D characterization is absolutely essential to experimentally prove the anomalous behavior predicted by our analysis with a simplified waveform as well as with the experimental beat patterns. Although we have performed a few experiments using beads of diameters of 1, 2 and 3 *μ*m (see Table S1), we are unable to verify the validity of the predicted anomalous trend because our 2D microscopy does not allow us to distinguish between symmetric and asymmetric bead-axoneme attachment. Future experiments with a multi-plane phase contrast microscope ^51^ is a promising step to resolve the full 3D geometry.

The design and fabrication of synthetic micro-swimmers is a challenging task in the growing field of smart drug delivery, and has recently become a multidisciplinary effort involving physicists, biologists, chemists and materials scientists. Flagella isolated from biological micro-organisms show a variety of waveforms and are promising candidates to provide a reliable power source for motility by an effective conversion of chemical energy from ATP hydrolysis into mechanical work. The *C. reinhardtii* flagellar-propelled micro-swimmers investigated here have the unique advantage that the beat frequency can be controlled via illumination, as we have described in our previous work ^42^, and chemical stimuli (e.g. calcium ions) can be used to trigger a transition from a circular to a straight swimming trajectories ^34,59^. They also serve as an ideal model-swimmer to investigate both experimentally and theoretically the contribution of different wave components in cargo propulsion dynamics. Our theoretical analysis as well as numerical simulations reveal the existence of an anomalous cargo transport regime, where contrary to expectation, the flagellar-propelled cargo swims faster as we increase the cargo size. This counter-intuitive behavior may play a crucial role in the design of future artificial flagellar-based propulsion systems. Finally, our analysis also highlights the contribution of the asymmetric cargo-flagellum attachment in the rotational velocity of the micro-swimmer. This turning mechanism should be also taken into account in manufacturing bio-inspired synthetic swimmers where a directional targeted motion is critical for delivery of drug-loaded cargoes.

## 4 Acknowledgment

The authors acknowledge J. Molacek, F. Nordsiek, B. Nasouri and unknown referees for valuable comments and S. Romanowsky, M. Müller and K. Gunkel for technical assistance. R.A. acknowledges support from the European Union’s Horizon 2020 research and innovation programme under grant agreement MAMI No. 766007. The authors acknowledge the Volkswagen Foundation Initiative LIFE (Project Living Foam) and the MaxSynBio Consortium, which is jointly funded by the Federal Ministry of Education and Research of Germany and the Max Planck Society. The authors also thank M. Lorenz and S. Bank at the Göttingen Algae Culture Collection (SAG) for providing the *C. reinhardtii* wild type strain SAG 11-32b.

## 5 Author contributions

A.G. designed the project. R.A. isolated flagella and performed the experiments. R.A., Y.S., S.G.P., A.P., E.B. and A.G. analyzed the flagella data. A.B. wrote the Matlab code to track the axonemes. A.G. and A.B. performed the theoretical analysis and simulations. All the authors discussed the results. A.G. wrote the first draft of the manuscript and A.P., E.B. and A.G. contributed to the revised version.

## 7 Supplemental Information

### 7.1 Rotational and translational velocities of an axoneme attached symmetrically to a bead

We used the simplified waveform given by Eq. 33 to calculate translational and rotational velocities of a freely swimming axoneme attached symmetrically from the basal end to a bead of dimensionless radius *r* = *R/L*. In the limit of small *C*_0_ = *κ*_0_*L/*(2*π*) and *C*_1_ = *κ*_1_*L/*(2*π*), we calculate rotational and translational velocities of the swimmer from Eq. 4 by calculating the propulsive forces and inverting the drag matrix **D** (see sections 2.3-2.3.1). We then take average over one beat cycle to obtain

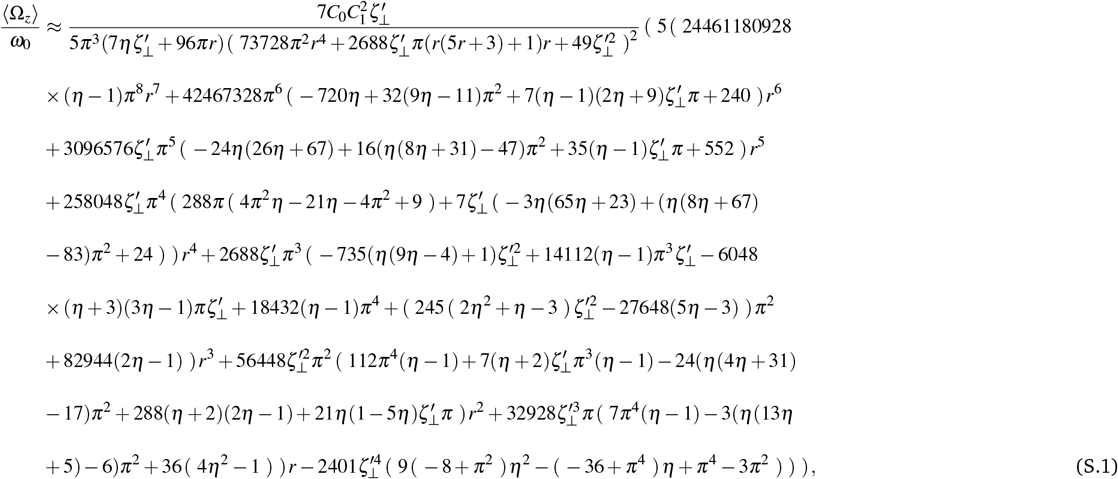

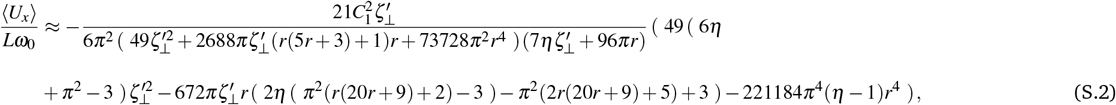

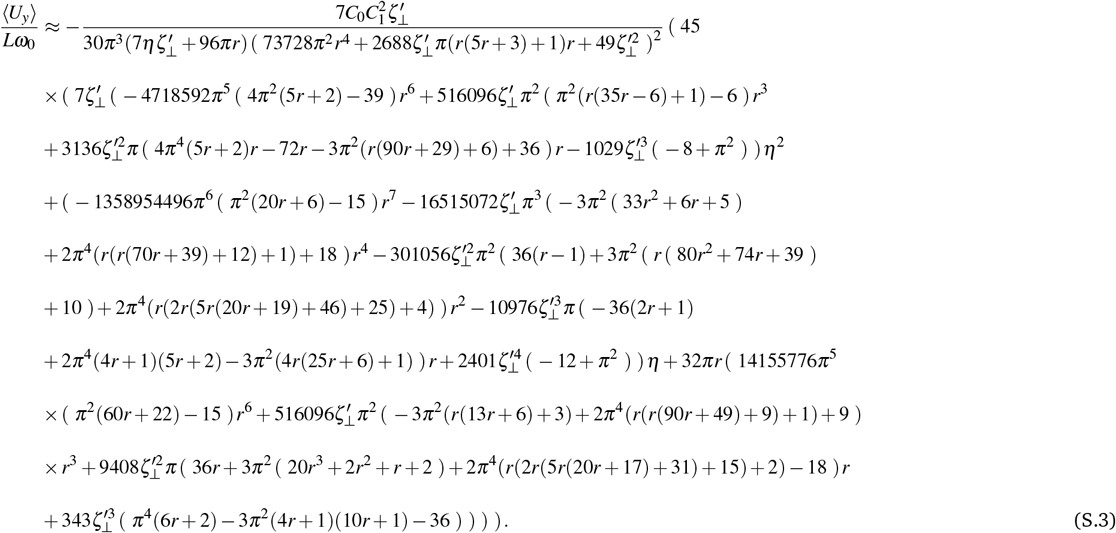

Here 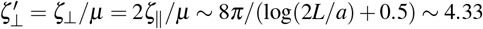 where *a* ~ 100 nm is the radius of axoneme, *L* ~ 10 *μ*m is the contour length of axoneme and we have assumed *η* = *ζ*_‖_ */ζ*_⊥_ = 0.5. Note that in the absence of the bead (*r* = 0) and with *η* = 0.5, Eqs. S.1-S.3 simplify to

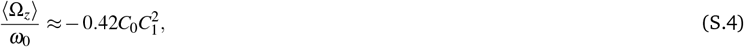

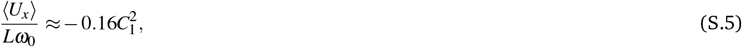

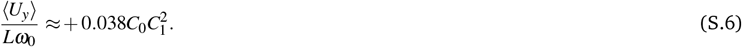

which are previously discussed in Refs. ^39,40^.

### 7.2 Rotational velocity of a bead attached asymmetrically to a freely-swimming axoneme

To mimic a sperm-like swimmer, we consider a model flagellum with only the main traveling wave component *C*_1_, setting the static component *C*_0_ to zero. We calculate the mean rotational velocity of an axoneme attached sideways to a bead using the matrix introduced in Eq. 30 and drag matrix of flagellum to obtain

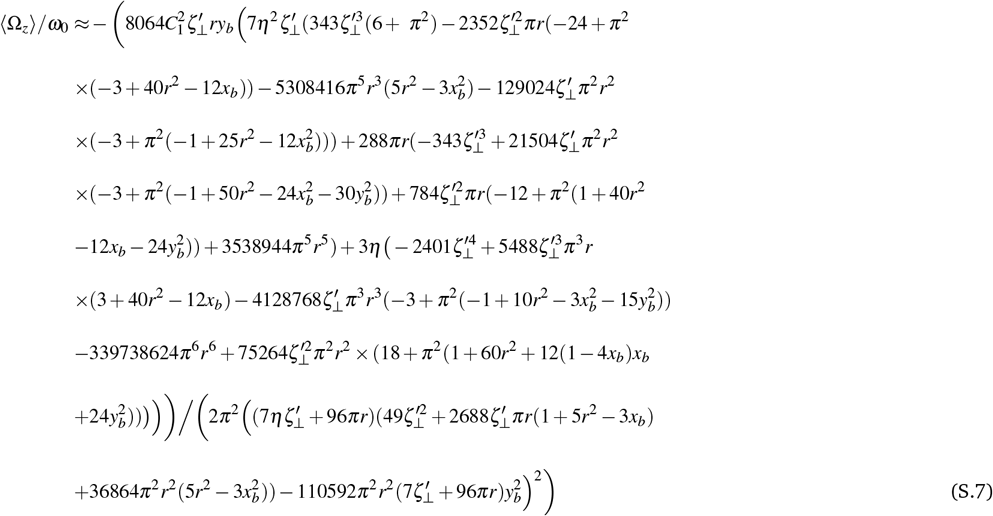

where 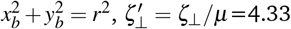 and *η* = *ζ*_‖_ */ζ*_⊥_=0.5.

### 7.3 Supplementary tables and figures

**Table S1.**
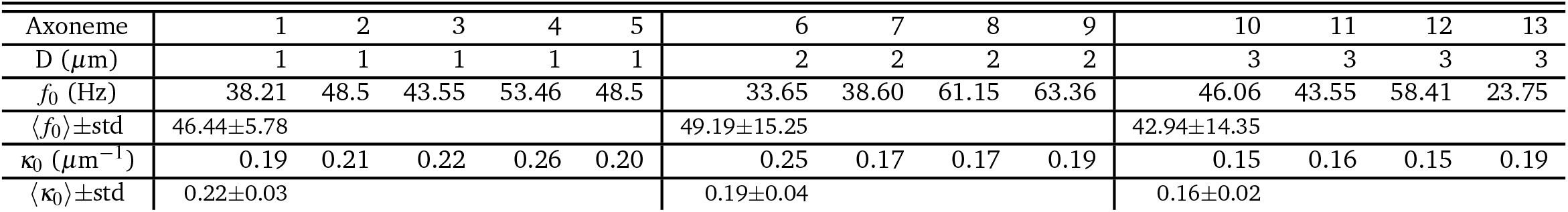
Frequencies and intrinsic curvature values of 12 different axonemes propelling beads of diameters *D*=1, 2 and 3 *μ*m. Mean and standard deviations are also presented. All axonemes are reactivated at [ATP]=200 *μ*M and [Ca^2+^]=0 mM. The axoneme No.1 corresponds to the example in Fig. 4A.

**Fig. S1.**
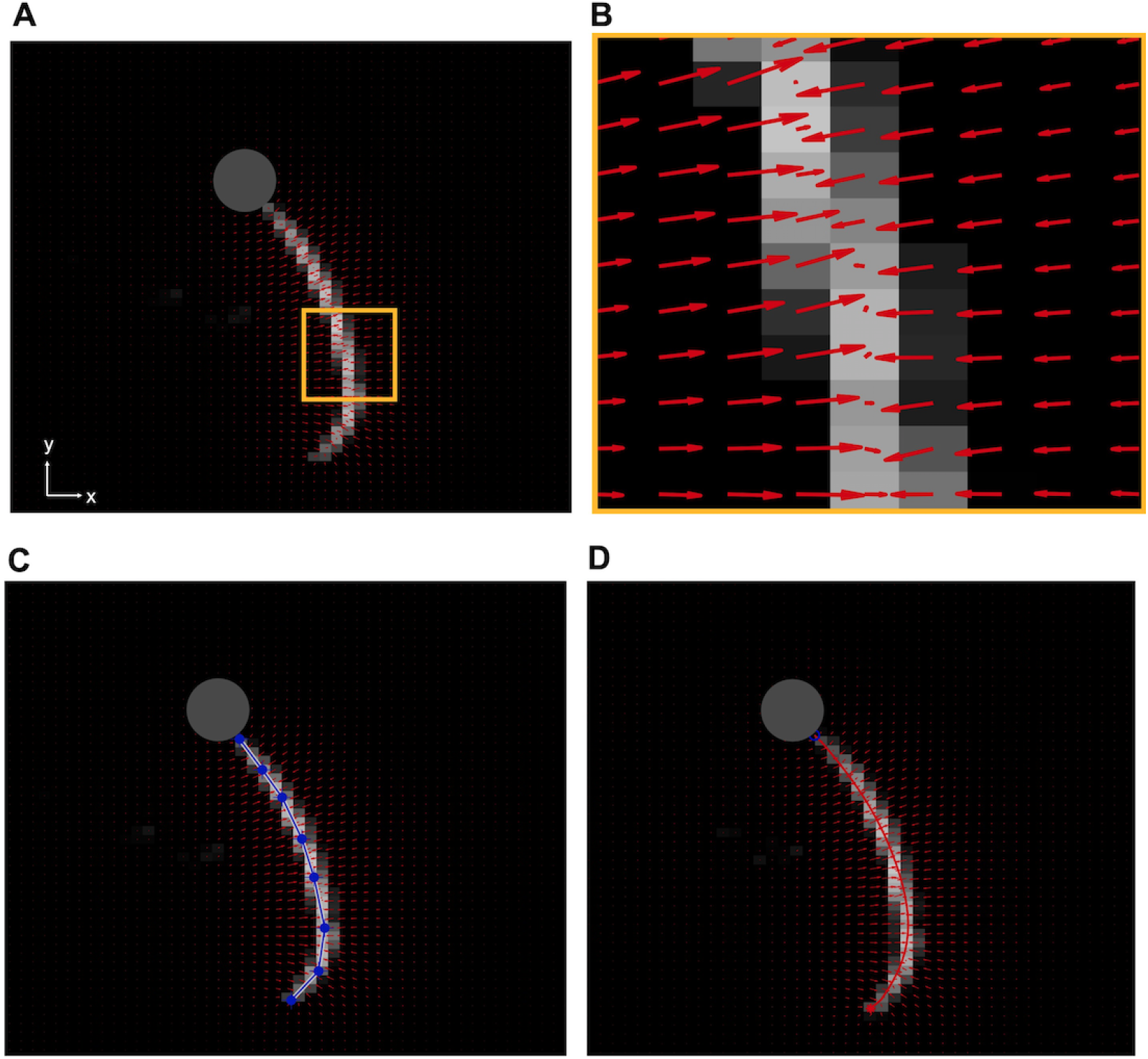
Gradient vector flow method implemented to track the flagellum. A) High intensity of the bead interferes with tracking, therefore it is removed by applying a threshold. B) The vector field calculated for small yellow region shown in panel A, converging toward pixels with maximum intensity. C) A polygonal line with eight nodes along the axonemal contour to initiate the snake. D) The tracking algorithm yields a discrete approximation of the axoneme’s contour represented by a set of *N* = 30 positions **r**(*s*_*i*_, *t*) = (*x*(*s*_*i*_, *t*), *y*(*s*_*i*_, *t*)), *i* = 1, …, *N*.

**Fig. S2.**
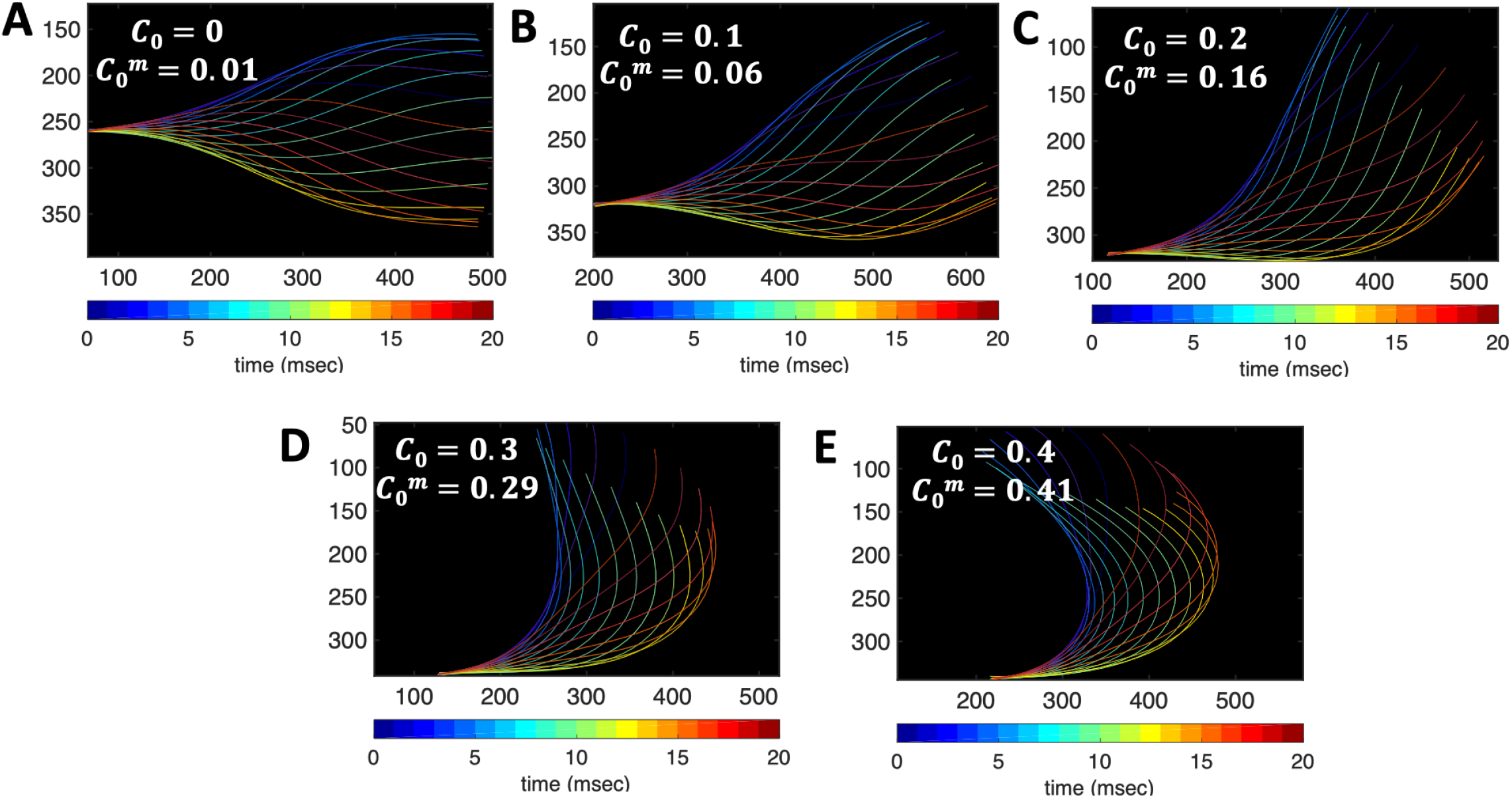
To measure the systematic error of our tracking GVF algorithm, we generated artificial data with known values of dimensionless mean curvature *C*_0_ = *κ*_0_*L/*(2*π*), and tracked the filaments using GVF. The measured values of mean curvatures of tracked filaments is given by 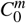. As it is shown in the panels A-E, the measured values deviate from the real values at small *C*_0_, but approaches the real values at higher *C*_0_. In other words, our algorithm has a smaller systematic error (less than 4%) for curved filaments with *C*_0_ *>* 0.3.

**Fig. S3.**
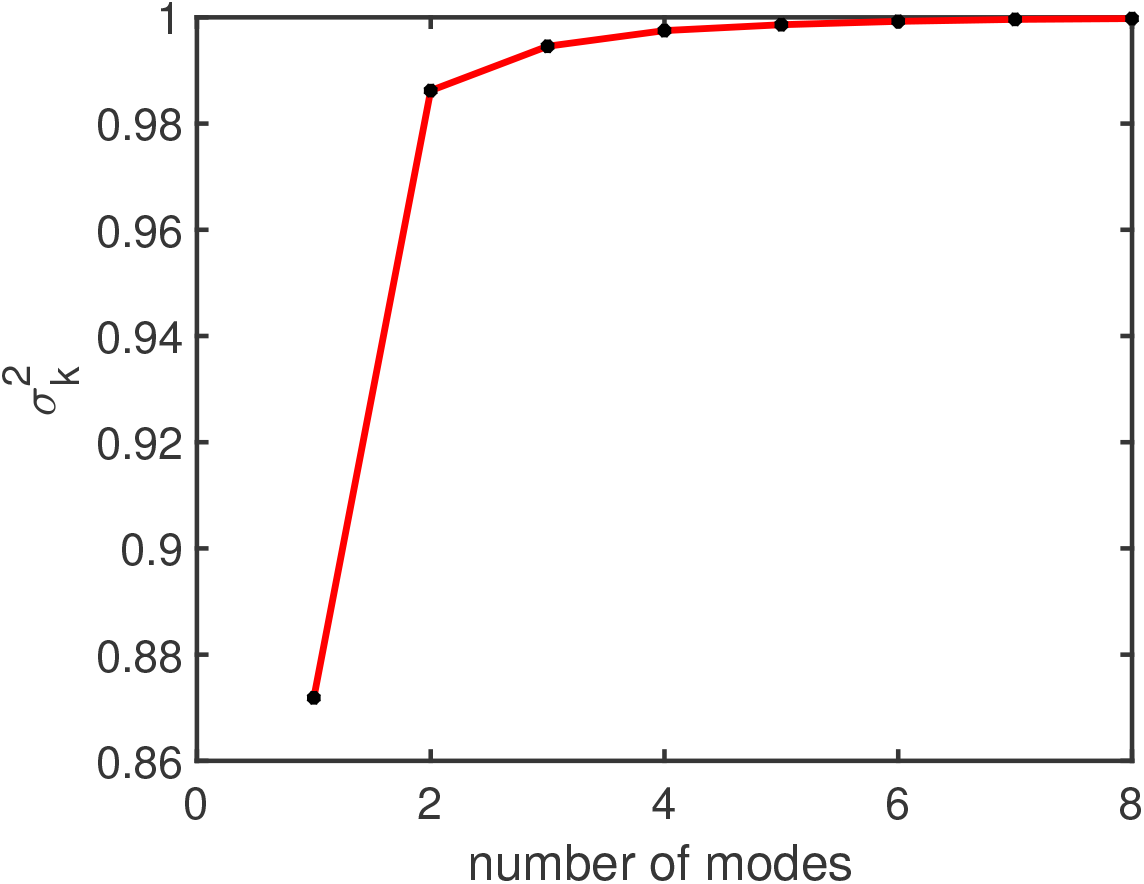
Fraction of the total variance 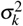, as defined in Sec. 2.4, plotted versus the number of PCA modes *n* for the axoneme shown in Fig. 4A.

**Fig. S4.**
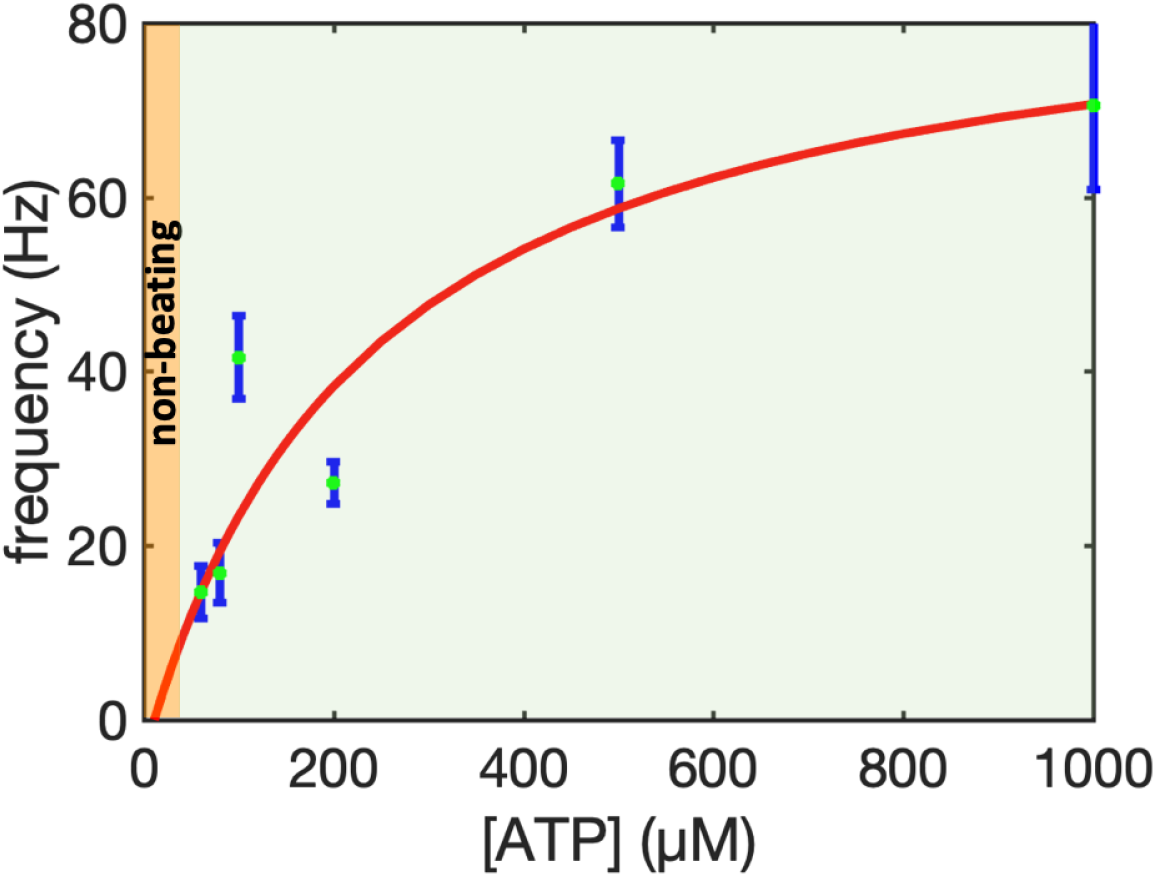
The frequency of reactivated axonemes depends on the [ATP] and follows a Michaelis-Menton type kinetics with a linear trend at small [ATP] and saturation at higher [ATP]. A minimum critical concentration of [ATP]_critical_=60 *μ*M is needed to reactivate the axonemes. In all the experiments [Ca^2+^]=0 mM.

**Fig. S5.**
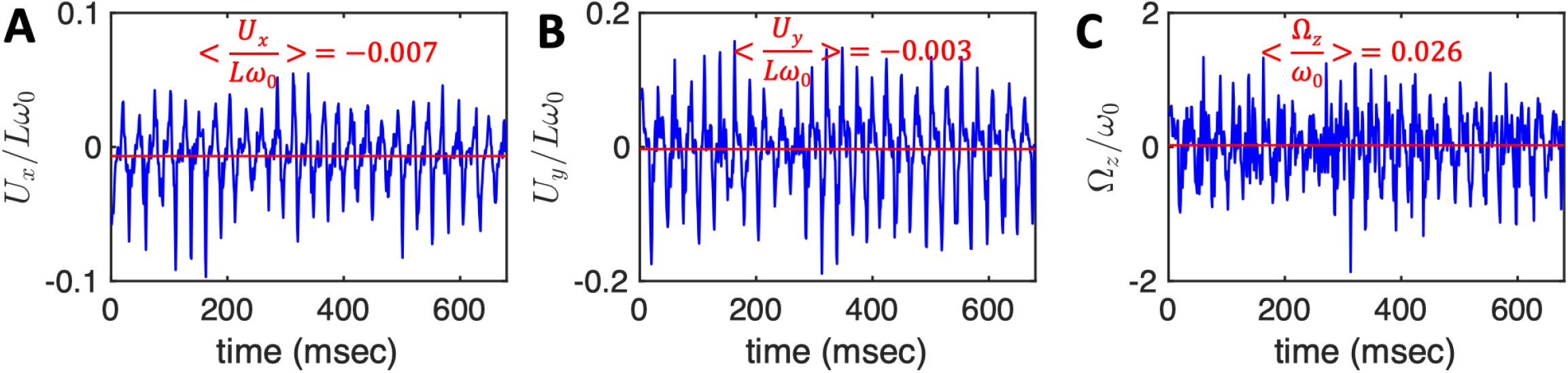
G) The translational and rotational velocities of the axoneme presented in Fig. 4A, measured in the swimmer-fixed frame, obtained with RFT simulations using the experimental beat pattern as input. The red lines mark the mean values.

**Fig. S6.**
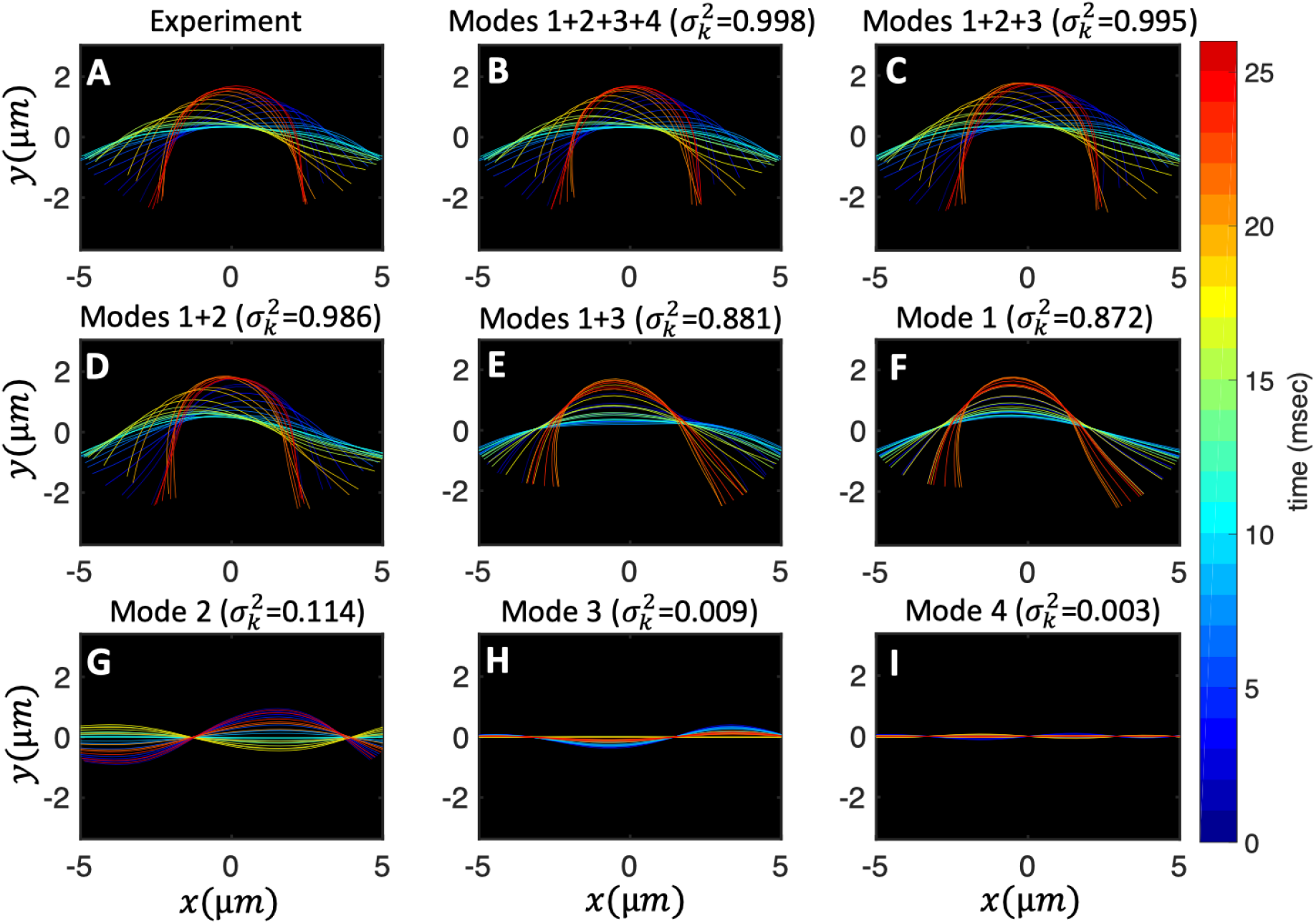
Experimental shapes over one beat cycle of the axoneme in Fig. 4A are compared with the axonemal shapes reconstructed using different combination of PCA modes as *θ* (*s, t*) = ⟨*θ* (*s, t*)⟩_*t*_ + Σ_*i*_*a*_*i*_(*t*)*M*_*i*_(*s*). For each panel, the fraction of the total variance 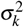 is also calculated; see section 2.4 for the definition of 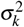. Note that shapes reconstructed using modes 2, 3 and 4 are approximately standing waves.

**Fig. S7.**
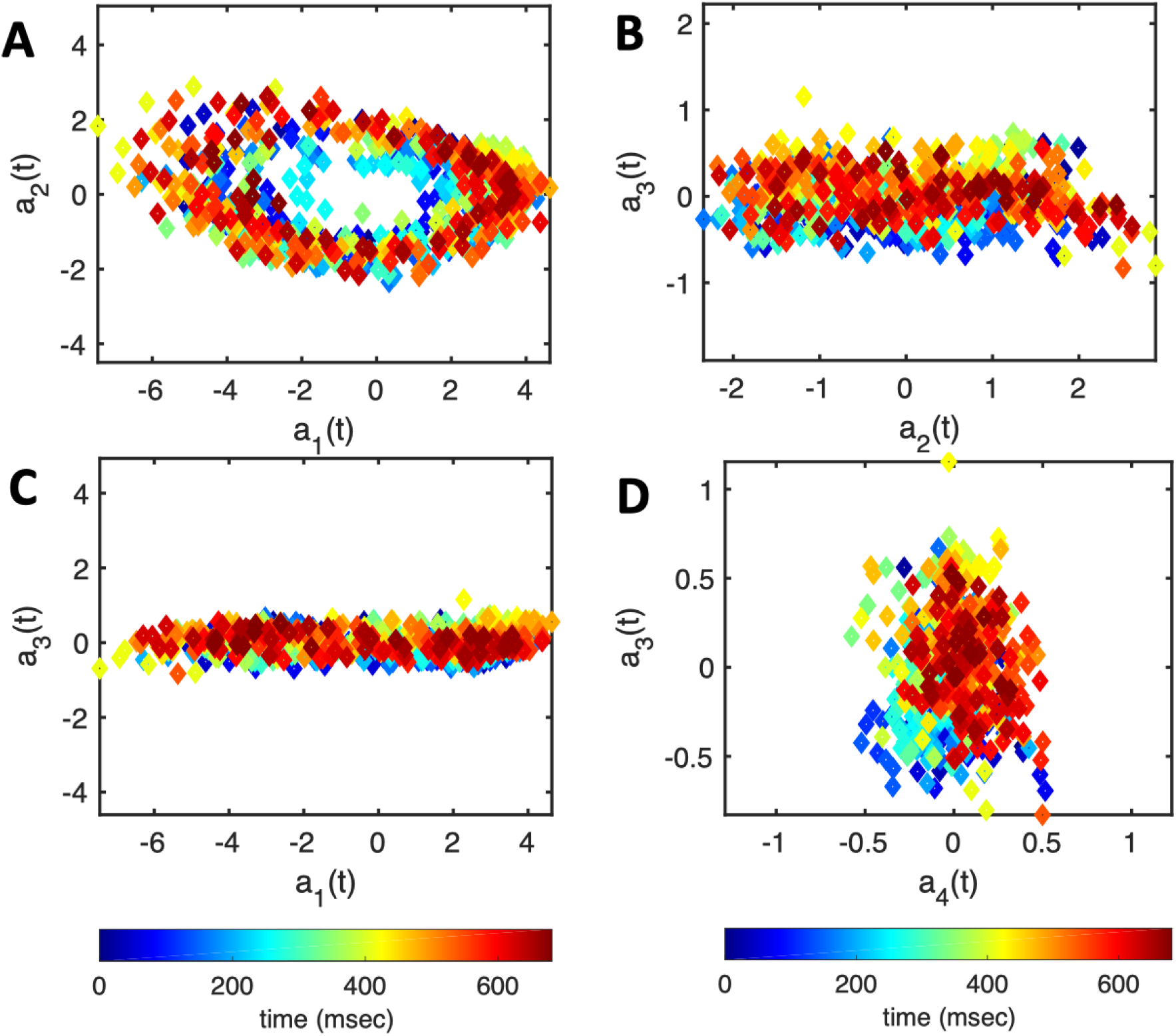
The time-dependent motion amplitudes *a*_1_(*t*) to *a*_4_(*t*) are plotted versus each other for the exemplary axoneme shown in Fig. 4A.

**Fig. S8.**
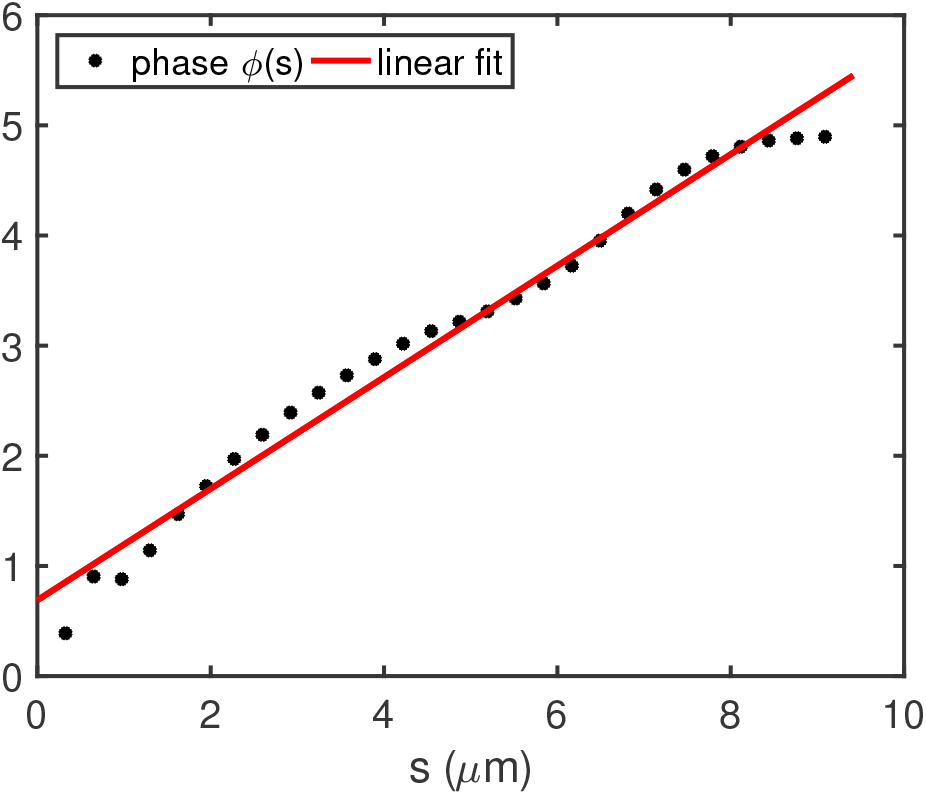
The phase *ϕ* (*s*) of traveling curvature waves of the axoneme presented in Fig. 4A, obtained by performing Fourier transform in time at each position *s* along the axonemal length. The wavelength is then calculated as *λ* = 2*πL/*(*ϕ* (*L*) −*ϕ* (0)) = 13.19 *μ*m.

**Fig. S9.**
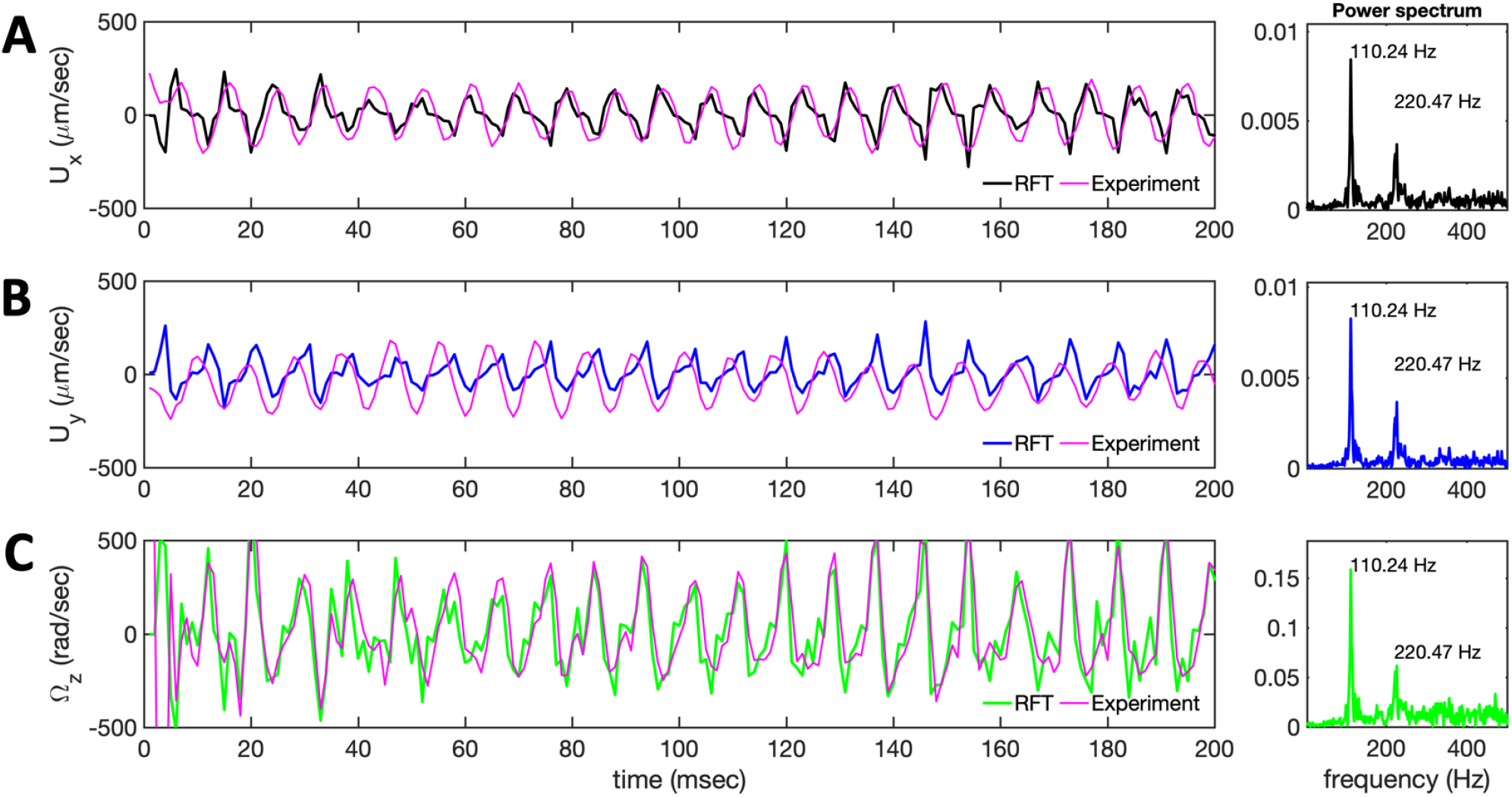
A-C) Velocity components of the bead’s center *U*_*x*_(*t*) and *U*_*y*_(*t*), and the rotational velocity of the bead Ω_*z*_(*t*) measured in the body-fixed frame of the exemplary axoneme in Video 7. Comparison with results obtained in the framework of resistive-force theory shows a good agreement. The power spectra show a dominant peak at the beat frequency of 110.24 Hz and its second harmonic. [ATP]=1 mM and [Ca^2+^]=0 mM.

**Fig. S10.**
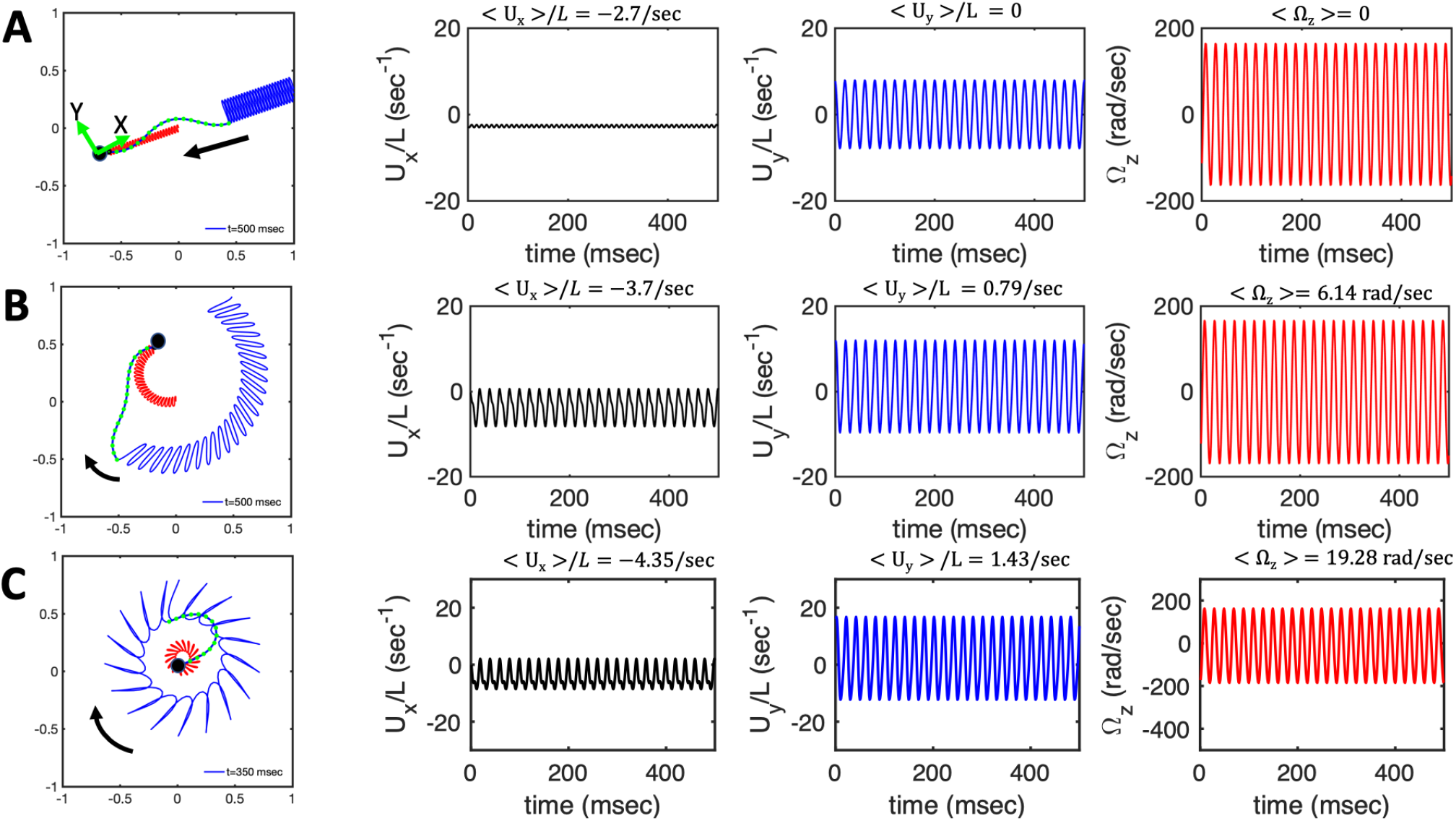
Simulations to show the effect of *C*_0_ at a fixed amplitude of dynamic mode *C*_1_ = 0.5. A bead of radius *R/L* = 0.1 is attached to the basal end. A-C) Swimming trajectory and mean translational and rotational velocities of the swimmer in the body-fixed frame. Parameters are *f*_0_ = 50 Hz, *η* = *ζ*_‖_ */ζ*_⊥_ = 0.5, A) *C*_0_ = 0, B) *C*_0_ = 0.25, and C) *C*_0_ = 0.5.

